# Synthesis and mechanical characterization of polyacrylamide (PAAm) hydrogels with different stiffnesses for large-batch cell culture applications

**DOI:** 10.1101/2024.09.17.613503

**Authors:** Charlotte Cresens, Samet Aytekin, Behrad Shaghaghi, Lotte Gerrits, Ana Montero-Calle, Rodrigo Barderas, Paul Kouwer, Susana Rocha

## Abstract

The impact of mechanical cues on cell behavior is increasingly being recognized, rendering hydrogel platforms that mimic the extracellular matrix indispensable in *in vitro* cell biology research. Here, we present a step-by-step protocol for synthesis and rheological characterization of polyacrylamide (PAAm) hydrogels with varying stiffnesses, produced as large circular unattached gels customizable in shape and size. We outline methods for their use in cell culture and downstream applications involving secretome or cell analysis, and protein visualization by fluorescence microscopy.

This protocol is based on the recent work of Shi & Janmey who describe a novel and straightforward method for the production of large PAAm hydrogels for bulk cell culture and mechanobiology studies.^1^ Their procedure results in one large gel that is not attached to a supporting surface and therefore can be transferred and/or stamped to generate PAAm gels of custom shapes and sizes. The aim of this step-by-step procedure is therefore not to improve the reported protocol, but to create a clearly outlined and repeatable protocol that enables a smooth implementation in any lab for a diverse audience. In addition, our protocol describes besides harvesting of cells also the collection of secretome for downstream biochemical analyses, as well as immunofluorescence labeling using antibodies that can readily be multiplexed for optimization of labeling conditions.

**Graphical abstract:** 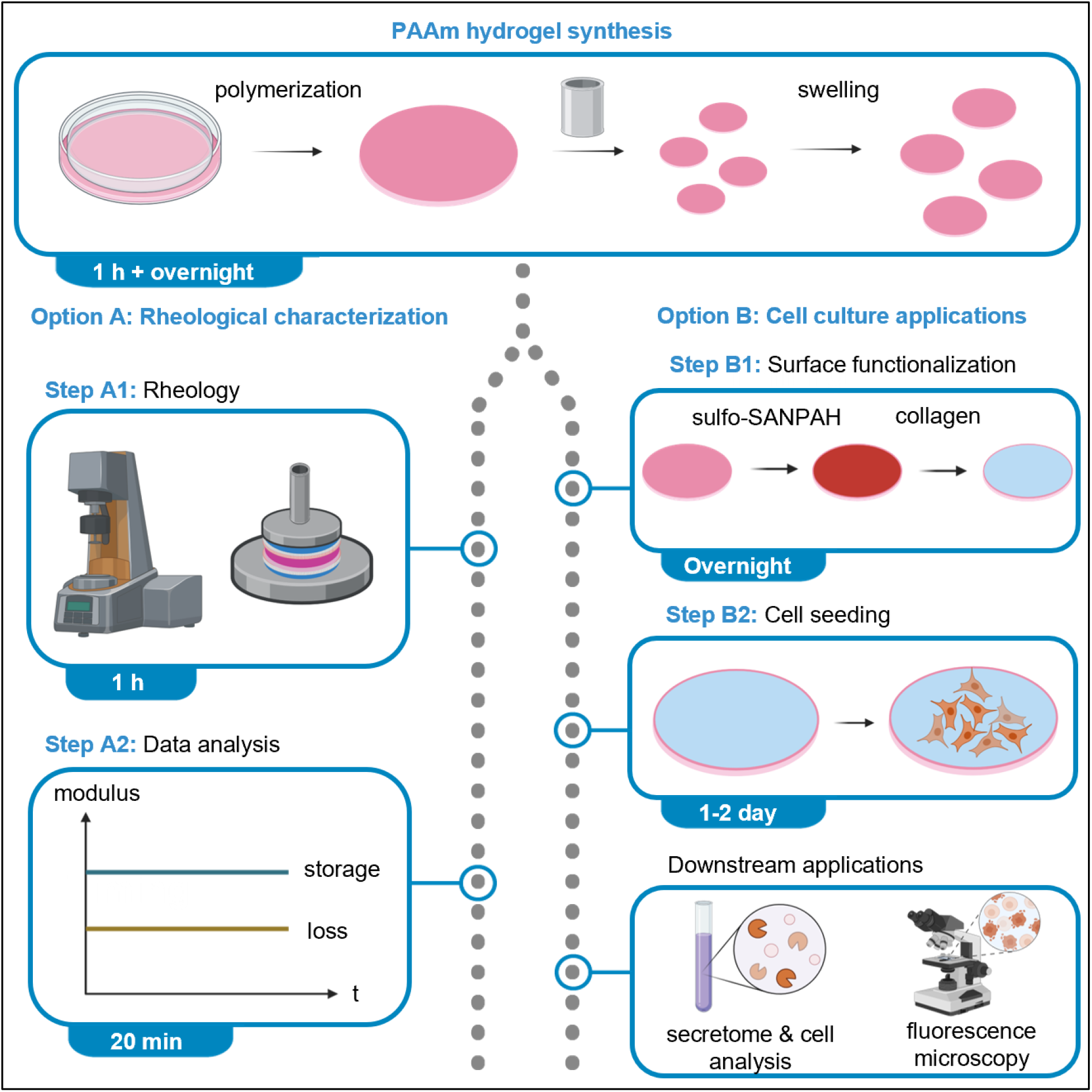

## Introduction

Since their implementation in 1997, polyacrylamide (PAAm, also referred to as PAA) hydrogels are widely being used as a versatile, inexpensive, and convenient platform for a wide variety of cell culture applications. They consist of acrylamide (AAm) monomer and N,N’-methylenebisacrylamide (MBAA) crosslinker, chemically crosslinked through a radical polymerization reaction catalyzed by ammonium persulfate (APS) and N,N,N’,N’-tetramethylethylenediamine (TEMED), where APS provides free radicals. The stiffnesses of these synthetic gels lie within the physiological range and can be precisely controlled by varying concentrations of AAm monomer and MBAA crosslinker, making them especially apt for mechanobiology studies where the influence of extracellular matrix (ECM) stiffness on cell behavior can be investigated. PAAm gels have been applied extensively to evaluate the effect of substrate stiffness on cell morphology, proliferation, adhesion and migration, molecular pathways, cell communication, differentiation, …^2–5^ A major advantage of PAAm gels is that they are biochemically inert and have to be functionalized with biological adhesion ligands, allowing for full control over the presented adhesion ligands, independently of the gels’ mechanical properties.^2,6,7^ In addition, PAAm gels are optically transparent at sufficiently low crosslinker concentrations (MBAA ≤ 0.5%), enabling their use in microscopy applications.^6^

Despite their numerous advantages, there are still some challenges or limitations associated to the use of PAAm hydrogels. Due to the small pore size and lack of sites for cellular adhesion, these hydrogels cannot be applied for 3D cultures where cells are embedded inside the hydrogel. Studies where cells are seeded on top of a functionalized 2D PAAm hydrogel, or even where cells are sandwiched between two PAAm hydrogel layers, have been reported.^8^ Notably, the reaction must be performed in between two surfaces to prevent oxygen from interfering with the radical polymerization. This also renders a flat surface that enables uniform cell attachment and growth.^3^ While immobilization of the hydrogel by covalent attachment to a surface is a common strategy, this is not a requirement, as demonstrated by Shi & Janmey who produce unattached PAAm gels by polymerization between two polystyrene surfaces.^1^

Despite – or because of – the widespread use of PAAm gels, large variations exist between stiffnesses reported for the same PAAm gel composition. On the one hand, differences might be introduced during the manufacturing when certain critical parameters are disparate, namely gelation temperature, reagent or PAAm gel storage time and temperature, mixing technique, …^3,5,9^ On the other hand, differences might be due to the highly variable measurement techniques used to characterize PAAm stiffness, ranging from nano-to microscale.^5^ We therefore propose this standardized protocol, where the steps involved in gel production, microscale rheology characterization, and functionalization for 2D cell culture are detailed. In addition, we describe two common downstream applications for cells grown on top of PAAm gels, namely harvesting of cells or medium for downstream biochemical analyses, and visualization of cell morphology or protein organization via fluorescence microscopy.

### Institutional permissions

Due to the acute toxicity, flammability, risk to eyes and skin, and long-term health hazards of several reagents used during PAAm gel production, some solutions and the PAAm gels should be prepared inside a Chemical Level 2 laboratory’s fume hood. Cell lines are handled exclusively in a Biosafety Level 2 laboratory and their use was approved by the ethical committee of UZ Leuven (reference number: S64861). Experiments are only performed after approval of a risk assessment and notification to the supervisor and HSE Department. Before adopting this protocol, make sure to comply with your institutional and national health, safety, and ethics regulations.

## Recipe 1: Prepare (sterile) DPBS 1x

### Timing: 10 min preparation + 4 h autoclaving

1. Dilute Dulbecco’s phosphate-buffered saline (DPBS) 10x without calcium or magnesium 1:10 in distilled water (dH_2_O) to obtain DPBS 1x.
  a. Add 50 mL DPBS 10x to 450 mL dH_2_O.
2. Optional for preparing sterile DPBS 1x: autoclave DPBS 1x in a heat-resistant container with the cap slightly open using a wet-cycle autoclave program. After the program is finished, close the container and cool to room temperature before use.

## Recipe 2: Prepare sterile dH_2_O

### Timing: 10 min preparation + 4 h autoclaving

3. Autoclave dH_2_O in a heat-resistant container with the cap slightly open using a wet-cycle autoclave program. After the program is finished, close the container and cool to room temperature before use.

## Recipe 3: Prepare APS 10% w/v solution

### Timing: 20 min

**NOTE:** Work in a chemical fume hood.

4. Prepare a 10% w/v ammonium persulfate (APS) solution by dissolving APS powder in dH_2_O.
  a. Weight 1 g APS powder in a 15 mL conical plastic tube.
  b. Add 10 mL dH_2_O.
  c. Dissolve by vortexing several minutes until the APS powder is totally dissolved, perform visual check for this.
5. Divide into 1 mL aliquots.
6. Store aliquots at -20 °C. Thaw immediately prior to use, avoid freeze-thawing.

## Recipe 4: Prepare sulfo-SANPAH 100 mg/mL solution

### Timing: 30 min

**NOTE:** Work in a sterile environment.

**CRITICAL:** Since sulfo-SANPAH is light-sensitive, it is important to protect the reagent from light.

7. Prepare a 100 mg/mL sulfosuccinimidyl 6-(4-azido-2-nitrophenylamino)hexanoate (sulfo-SANPAH) solution by dissolving sulfo-SANPAH powder in anhydrous dimethylsulfoxide (DMSO).
  a. To a vial containing 50 mg sulfo-SANPAH, add 200 µL anhydrous DMSO.
  b. Dissolve the powder by pipetting and transfer everything to a sterile 1.5 mL Eppendorf tube.
  c. Rinse the original vial 3x with 200 µL, 50 µL and 50 µL anhydrous DMSO, and transfer everything to the same Eppendorf tube. **TIP:** After each rinsing step the solution becomes less dark-red.
  d. Ensure a homogenous solution by pipetting the sulfo-SANPAH solution in the 1.5 mL Eppendorf tube.
  **CRITICAL**: Sulfo-SANPAH is sensitive to moisture, so only employ anhydrous DMSO to preserve reagent integrity.
8. Divide into 25 µL aliquots.
9. Store aliquots at -20 °C protected from light. Thaw immediately prior to use, avoid freeze-thawing.

## Recipe 5: Prepare NaOH 1 M solution

### Timing: 30 min preparation + 1 h dissolving

**NOTE:** Work in a chemical fume hood.

10. Prepare a 1 M sodium hydroxide (NaOH) solution by dissolving NaOH tablets in dH_2_O.
  a. Weight 40 g NaOH.
  b. Place a glass beaker containing a magnetic bar on a stirring plate and add 50 mL dH_2_O.
  c. Add approximately 5 NaOH pellets at a time and wait a few minutes until they are fully dissolved before adding more. It may take up to 1 hour until all NaOH is dissolved. **CAUTION**: The reaction is extremely exothermic.
  d. Transfer the volume to a volumetric flask of 250 mL.
  e. Rinse the beaker 3x with 50 mL dH_2_O for every step and transfer everything to the same volumetric flask.
  f. Complete the volume to 250 mL using dH_2_O.
11. Store the solution at room temperature in a well-ventilated area.

## Recipe 6: Prepare sterile AA 0.5% solution with pH 3.40

### Timing: 30 min

**NOTE:** Work in a chemical fume hood.

12. Prepare a 0.5% acetic acid (AA) solution by diluting 100% AA in dH_2_O.
  a. To a glass beaker with stirring magnet, add 298.5 mL dH_2_O and 1.5 mL acetic acid 100%.
  b. Homogenize the solution by stirring.
13. Adjust the pH to 3.40 using NaOH 1 M solution (prepared using Recipe 5).
  a. Calibrate the pH meter according to the manufacturers’ instructions.
  b. Dip the pH electrode into the stirring acetic acid solution.
  c. Add dropwise the 1 M NaOH solution while monitoring the pH, until it reaches 3.40. **TIP:** For a volume of 300 mL, approximately 800 – 1000 µL 1 M NaOH are required.
14. Sterilize the solution into 50 mL aliquots using a 50 mL syringe and syringe filter (0.2 µm pore size).
15. Store the solution at room temperature.

## Recipe 7: Prepare sterile cell culture and starvation medium

### Timing: 15 min

**NOTE:** Work in a sterile environment.

**NOTE:** Sterile starvation medium is optional for secretome and cell harvesting to avoid critical interference of FBS in subsequent analysis (Downstream application 1).

16. Prepare aliquots of cell culture additives for easier handling.
  a. Aliquot fetal bovine serum (FBS) to 50 mL, aliquot GlutaMAX™ to 5 mL, aliquot gentamycin to 500 µL.
17. Prepare culture medium suitable for culture of KM12L4a cells: Dulbecco’s modified eagle medium (DMEM) supplemented with 10% FBS, 1% GlutaMAX™ and 0.1% gentamycin sulphate.
  a. For preparing 500 mL medium, remove 55.5 mL DMEM from the 500 mL flask. Add 50 mL FBS, 5 mL GlutaMAX™ and 500 µL gentamycin sulphate. **TIP:** Notice that a certain volume DMEM is removed before adding the supplements to obtain the exact concentrations of FBS, GlutaMAX™ and gentamycin as indicated. Failing to remove the DMEM at the start, will result in slightly different final concentrations. **CRITICAL:** The composition of the cell culture medium may need to be adjusted, depending on the type of cell line used.
18. *Optional:* Prepare starvation medium, which is standard cell culture medium without FBS.
  a. For preparing 500 mL starvation medium, remove 5.5 mL DMEM from the 500 mL flask. Add 5 mL GlutaMAX™ and 500 µL gentamycin.
19. Store media at 4 °C. Store leftover aliquots of FBS, GlutaMAX™ and gentamycin at -20 °C.

## Recipe 8: Prepare sterile EDTA solution

### Timing: 15 min

**NOTE:** Work in a sterile environment.

For general cell harvesting, prepare a trypsin-EDTA 1x solution in DPBS 1x.

20. For example, add 5 mL of trypsin-EDTA 0.5% solution (10x concentrated) to 45 mL of DPBS 1x (prepared using Recipe 1).
21. Store the solution at 4 °C.

For proteomics cell harvesting (Downstream application 1), prepare a sterile 4 mM EDTA solution in commercial DPBS 1x.

22. Sterilize a portion of the 0.5 M ethylenediaminetetraacetic acid (EDTA) stock solution using a 15 mL syringe and syringe filter (0.2 µm pore size).
23. Prepare a 4 mM EDTA solution in commercial DPBS 1x.
  a. Add 400 µL sterile 0.5 M EDTA stock solution to 49.6 mL commercial DPBS 1x.
24. Store the solution at 4 °C.

## Recipe 9: Prepare PFA 4% solution

### Timing: 10 min

**NOTE:** Work in a chemical fume hood.

**NOTE:** This is optional for immunolabeling (Downstream application 2).

25. Prepare a 4% paraformaldehyde (PFA) solution in DBPS 1x.
  a. Add the content of 1 vial PFA 16% (10 mL) to 30 mL DPBS 1x (prepared using Recipe 1).
26. Store the solution at 4° C.

## Recipe 10: Prepare Triton X-100 0.1% solution

### Timing: 15 min

**NOTE:** Work in a chemical fume hood.

**NOTE:** This is optional for immunolabeling (Downstream application 2).

27. Prepare a 10% Triton X-100 solution in DPBS 1x (prepared using Recipe 1).
  a. Add 45 mL DPBS 1x in a glass beaker containing a magnetic bar on a stirring plate.
  b. Add 5 mL Triton X-100 100%. **NOTE:** Triton X-100 is very viscous.
  c. Stir until fully dissolved.
28. Dilute the solution prepared in the previous step to a 0.1% Triton X-100 solution using DPBS 1x (prepared using Recipe 1).
  a. Add 500 µL Triton X-100 10% solution in 49.5 mL DBPS 1x.
29. Store both solutions at 4 °C.

## Recipe 11: Prepare blocking buffer

### Timing: 10 min

**NOTE:** This is optional for immunolabeling (Downstream application 2).

30. Prepare blocking buffer containing 10% FBS in DPBS 1x (prepared using Recipe 1).
  a. Add 1 mL FBS 100% to 9 mL DPBS 1x.
31. Store the solution at 4 °C for maximally 2 days.

## Recipe 12: Prepare DAPI 5 mg/mL solution

### Timing: 10 min

**NOTE:** Work in a chemical fume hood and protect DAPI from exposure to light.

**NOTE:** This is optional for immunolabeling (Downstream application 2).

32. Prepare a 5 mg/mL 4’,6-Diamidino-2-Phenylindole (DAPI) solution by dissolving DAPI powder in dH_2_O.
  a. To a vial containing 10 mg DAPI powder, add first 1 mL dH_2_O. Dissolve the powder by pipetting and transfer everything to a sterile 15 mL conical tube.
  b. Rinse the original vial with 500 µL, 250 µL and 250 µL dH_2_O, and transfer everything to the same conical tube. **TIP:** After each rinsing step the solution becomes less yellow.
  c. Ensure a homogenous solution by pipetting up and down the solution in the conical tube
33. Divide into 25 µL aliquots.
34. Store aliquots at -20 °C protected from light. Thaw immediately prior to use, avoid freeze-thawing.

## Key resources table

| REAGENT or RESOURCE | SOURCE | IDENTIFIER |
| --- | --- | --- |
| **Antibodies** |  |  |
| Optional for immunolabeling (Downstream application 2):<br>Monoclonal mouse primary antibody against human, mouse, rat alpha tubulin [DM1A]<br>Store at -20 °C for long-term storage. Avoid freeze-thaw cycles. | Abcam | ab7291 |
| Optional for immunolabeling (Downstream application 2):<br>Goat-anti-mouse (GAM) Atto647N secondary antibody<br>Store at -20 °C protected from light for long-term storage. Avoid freeze-thaw cycles. | Sigma-Aldrich (Merck) | 50185 |
| **Bacterial and virus strains** |  |  |
| NA | NA | NA |
| **Biological samples** |  |  |
| NA | NA | NA |
| **Chemicals, peptides, and recombinant proteins** |  |  |
| Dulbecco's phosphate-buffered saline (DPBS)<br>10X, no calcium, no magnesium. | Gibco (Thermo Fisher Scientific) | 14200075 |
| Optional for cell harvesting (Downstream application 1):<br>Dulbecco's phosphate-buffered saline (DPBS)<br>1X, no calcium, no magnesium. | Gibco (Thermo Fisher Scientific) | 14190144 |
| Acrylamide (AAm) solution<br>40% w/v solution.<br>Mr = 71.08 g/mol.<br>Store at 2-8 °C.<br>**CAUTION:** Is acutely toxic upon ingestion/inhalation, and represents a long-term health hazard. | Sigma-Aldrich (Merck) | A4058 |
| N,N'-nethylenebisacrylamide (MBAA) solution<br>• 2% w/v solution<br>• Mr = 154.17 g/mol.<br>• Store at 2-8 °C.<br>• **CAUTION:** Represents a long-term health hazard. | Sigma-Aldrich (Merck) | M1533 |
| Ammonium persulfate (APS) powder<br>• Mr = 228.18 g/mol.<br>• Store powder at room temperature in a dry, cool and well-ventilated location.<br>• **CAUTION:** Is hazardous upon ingestion/inhalation, causes severe skin burns, is oxidizing and represents a long-term health hazard. | Bio-Rad | #1610700EDU |
| N,N,N',N'-tetramethylethylenediamine (TEMED) solution<br>• ≥99% solution.<br>• Mr = 116.24 g/mol.<br>• Store at room temperature in a dry and well-ventilated location.<br>• **CAUTION:** Is acutely toxic upon ingestion/inhalation, causes severe skin burns and is highly flammable. It has been associated to male infertility. | Sigma-Aldrich (Merck) | T7024 |
| Anhydrous dimethyl sulfoxide (DMSO)<br>• Mr = 78.13 g/mol.<br>• Store at room temperature.<br>• **CAUTION:** Is highly flammable. | Invitrogen (Thermo Fisher Scientific) | D12345 |
| Sulfosuccinimidyl 6-(4-azido-2-nitrophenylamino)hexanoate (sulfo-SANPAH) powder<br>• Mr = 492.4 g/mol.<br>• Store at -20 °C protected from light, in a dry environment.<br>• **CAUTION:** Causes serious eye irritation. | Sigma-Aldrich (Merck) | 803332 |
| Acetic acid (AA) 99.8% solution<br>• Mr = 60.05 g/mol.<br>• Store at room temperature in a well-ventilated location.<br>• **CAUTION:** Flammable, causes severe skin burns and eye damage. | Acros Organics | 64-19-7 |
| Sodium hydroxide (NaOH) pellets<br>• Mr = 40.00 g/mol.<br>• **CAUTION:** Causes severe skin burns and eye damage. May be corrosive to metals. | AnalaR NORMAPUR (VWR) | 28244.295 |
| Collagen type I from rat tail<br>• 3 mg/mL liquid solution.<br>• Store at 4 °C protected from light. | Thermo Fisher Scientific | A1048301 |
| Dulbecco's Modified Eagle Medium (DMEM)<br>• high glucose, no glutamine, no phenol red.<br>• Store at 4 °C protected from light. | Gibco (Thermo Fisher Scientific) | 31053028 |
| Fetal Bovine Serum (FBS)<br>• Store at -20 °C. | Sigma-Aldrich (Merck) | F7524 |
| GlutaMAX™ Supplement<br>Manufacturer reports stability across a wide range of temperatures. In our lab, GlutaMAX™ is stored at -20 °C. | Gibco (Thermo Fisher Scientific) | 35050038 |
| Gentamycin sulphate solution<br>Store at -20 °C.<br>**CAUTION:** May cause an allergic skin reaction, cause allergy or asthma symptoms or breathing difficulties if inhaled. | Carl Roth | 2475.1 |
| Trypsin-ethylenediaminetetraacetic acid (Trypsin-EDTA) 0.5% solution, no phenol red<br>Is a 10x solution.<br>Store at -20 °C.<br>**CAUTION:** Represents a long-term health hazard. | Thermo Fisher Scientific | 15400054 |
| Optional for proteomics cell harvesting (Downstream application 1):<br>Ethylenediaminetetraacetic acid (EDTA)<br>0.5 M solution.<br>Store at +20 °C.<br>**CAUTION:** Causes serious eye irritation and represents a long-term health hazard. | Thermo Fisher Scientific | J15694.AE |
| Trypan blue 0.4% solution<br>Store at 4 °C.<br>**CAUTION:** Represents a long-term health hazard. | VWR | K940 |
| Optional for immunolabeling (Downstream application 2):<br>Paraformaldehyde (PFA) 16% solution, methanol-free<br>Store at room temperature in a dry, cool, and well-ventilated location.<br>**CAUTION:** Causes serious eye irritation, harmful when swallowed or inhaled. May cause cancer and respiratory irritation. | Thermo Fisher Scientific | 28908 |
| Optional for immunolabeling (Downstream application 2):<br>Triton X-100 detergent, molecular biology grade<br>Store at room temperature in a dry, cool, and well-ventilated location.<br>**CAUTION:** Acutely toxic, causes serious eye damage and skin irritation. Very toxic to aquatic life. | Sigma-Aldrich (Merck) | T8787 |
| Optional for immunolabeling (Downstream application 2):<br>4',6-Diamidino-2-Phenylindole (DAPI)<br>Store at -20 °C protected from light.<br>**CAUTION:** May cause genetic defects. | Invitrogen (Thermo Fisher Scientific) | D1306 |
| **Critical commercial assays** |  |  |
| NA | NA | NA |
| **Deposited data** |  |  |
| Measurement details of PAAm hydrogel library characterization are shown in Table S1. Rheology Trios data files are available upon reasonable request. |  |  |
| **Experimental models: Cell lines** |  |  |
| KM12L4a cell line | Fidler's lab (MD Anderson Cancer Center) | RRID:CVCL_W220 |
| **Experimental models: Organisms/strains** |  |  |
| NA | NA | NA |
| **Oligonucleotides** |  |  |
| NA | NA | NA |
| **Recombinant DNA** |  |  |
| NA | NA | NA |
| **Software and algorithms** |  |  |
| Trios software for the rotational rheometer | TA Instruments | Version 5.5.0.323 |
| CytoSMART™ App software | Corning | ND |
| Microsoft excel | Office | Version 2302 |
| Optional for immunolabeling (Downstream application 2):<br>Las X 3D Viewer | Leica Microsystems GmbH | ND |
| **Other** |  |  |
| Petri dish, 92 x 16 mm, transparent, with ventilation cams<br>Made from polystyrene (PS).<br>Base dish: 87.2 mm diameter x 13.5 mm height.<br>Lid: 91.2 mm diameter x 7.5 mm height.<br>When the base dish with the open side up is placed inside the lid and centered with respect to it, the polymerization chamber has an open edge of approximately 1 mm on all sides. | Sarstedt | 82.1473.001 |
| Large square or circular Petri dish, transparent<br>Made from polystyrene (PS).<br>Large square Petri dish can be for example 119 mm length of one side x 15 mm height.<br>Large circular Petri dish can be for example 140 mm base diameter x 20 mm height. | For square: Corning<br>For circular: Merck | For square: BP124-05<br>For circular: Z717231 |
| Optional for immunolabeling (Downstream application 2):<br>Immunolabeling incubation chamber<br>See section "Materials and equipment" for more information. | ND | ND |
| Optional for immunolabeling (Downstream application 2):<br>6-well glass-bottom plate with high performance #1.5 cover glass | Cellvis | P06-1.5H-N |
| Corning® 28 mm Diameter Syringe Filters, 0.2 µm Pore SFCA Membrane, Sterile, Individually Packaged | Corning | 431219 |
| 50 mL sterilization syringe<br>50 mL BD® Eccentric Luer Slip Tip Syringe only. | BD | 300866 |
| 20 mL sterilization syringe<br>Inject® solo 20ML luer excentrisch. | Braun (Injekt) | 4606205V |
| Dust-free precision wipes | Kimtech | 7552 |
| Sandpaper: P400 Grit Extra Fine Sanding Sheet, 280mm x 230mm | 3M | P400 |
| Double-sided adhesive tape transparent, 25 mm x 50 m | Tesa | 64621 |
| Flat, round-tipped precision tweezer | Chemlab analytical | LL6272047 |
| Micro scissors | Carl Roth | EPA0.1 |
| Scoop or spatula with flat and thin surface<br>Optional to simplify handling of PAAm gels.<br>e.g. alginate spatula, but ideally with a bend between the blade and handle. | Saillart | HWN 272-22 |
| Stamp for creating gels of custom shapes and sizes<br>See section “Materials and equipment” for more information. | ND | ND |
| Multi-parameter analyzers used for pH measurement | Consort | C1010 |
| Branson Ultrasonic Cleaner used for sonication | Fisher Scientific | 2510E-DTH |
| Stress-controlled rotational rheometer | TA Instruments | Discovery HR-2 |
| Steel parallel plate geometry with 20 mm plate diameter (Stainless Steel, 20 mm Plate, NST) | DHR & AR-G2 | 511200.906 |
| Rocker and 3D shakers RS series, seesaw | Carl Roth (Phoenix Instrument) | RS-RS 5 |
| UV-lamp with 365 nm wavelength emission<br>See section “Materials and equipment” for more information. | Camag | 022.9120 |
| Cell culture humidified incubator at 37 °C and 5% CO<sub>2</sub> (e.g. Heracell™ VIOS 160i CO2 Incubator) | Thermo Fisher Scientific | 51033557 |
| Hettig Universal centrifuge | Hettig | 320R |
| Corning® Cell Counter | Corning | 6749 |
| Leica DM IL LED Inverted Laboratory Microscope | Leica Microsystems GmbH | ND |
| Optional for imaging of immunolabeled targets (Downstream application 2):<br>Leica TCS SP8 confocal laser scanning microscope<br>405/488/514/552/638 nm diode excitation lasers.<br>63x water immersion objective (Leica HC PL APO CS2 63x/1.20).<br>highly sensitive hybrid detectors (HyD SMD), and use a transmitted light detector for transmission images. | Leica Microsystems GmbH | ND |

## Materials and equipment

- All numerical values concerning the dimensions of PAAm gels or related materials represent their diameter, unless stated otherwise.
- Unless stated otherwise, DPBS 1x indicates the DPBS that is prepared using Recipe 1.
- AAm and MBAA are acutely toxic upon ingestion/inhalation, and represent a long-term health hazard. Using the pre-made solutions is therefore preferred over weighting and dissolving the powders. Note that these limit the final percentage of AAm in the PAAm recipes, and therefore the maximum PAAm gel stiffness that can be obtained (Table 1). In case higher AAm percentages are desired, weighting and dissolving the powders will be required.

**Table 1:** PAAm hydrogel library. Stiffness (Youngs moduli) and practical gel composition necessary for production of one large PAAm hydrogel are reported for the different gel compositions (in the different rows). Stiffnesses were measured and calculated by rheology as described in this protocol, average ± error taken over minimally 3 technical replicates are reported. Con final = final concentration. V/V final = final volume/volume.
- We used a circular polystyrene Petri dish of 92 × 16 mm as polymerization chamber in which we added 9.5 mL of the PAAm liquid complete mixture (see Key resources table and Major step 1). Petri dishes of other sizes can be used, the only requirement for creating the polymerization chamber is that the diameter of the Petri dish lid is larger than the diameter of the base.
- We used several different stamps for creating gels of custom shapes and sizes. In general, a tissue-culture stamp or patisserie garnish stamp with a sharp edge and custom-shape and -size can be used.
  ∘ For rheological characterization (Major option A), a circular 20 mm diameter tissue-culture stamp with sharp edge was used. This stamp was custom-made in stainless steel, but can alternatively be purchased (e.g. Rennsteig Werkzeuge 140 020 0 Holpijp 20 mm from Conrad, reference 1901241 - 8J)
  ∘ For cell culture applications (Major option B), a circular 75 mm diameter tissue-culture stamp with sharp edge was used. This stamp was custom-made, but can alternatively be purchased (e.g. Rennsteig Werkzeuge 140 075 0 Holpijp 75 mm from Conrad, reference 1901276 - 8J)
  ∘ For immunolabeling and visualization in fluorescence microscopy (Downstream application 2), a circular 10 mm diameter tissue-culture stamp with sharp edge was used. This stamp was custom-made, but can alternatively be purchased (e.g. Rennsteig Werkzeuge 140 010 0 Holpijp 10 mm from Conrad, reference 1901231 - 8J)
- For sulfo-SANPAH functionalization (Major option B), a UV-lamp with 365 nm wavelength emission was used. The protective plate was removed, and the lamp was set up in a closed protective box positioned approximately 6 cm above the sample. UV-power was measured to be 8.92 ± 0.14 mW for a rectangular area of 20.30 × 1.00 mm. For different powers or configurations, incubation times may need to be adjusted.
- Immunolabeling (Downstream application 2) was performed in a 12 mm chamber of 5 mm deep, and a recess of 4 mm that protrudes from the chamber and accommodates the round-tipped tweezers. However, any small container with a lid in which the gel can still be handled with the round-tipped tweezers will suffice. For example, a 24-well plate with plastic, flat bottom (Starstedt, reference 83.3922) is a commercially available alternative.

| Composition | Gel characterization | Gel composition |  |  |  | Practical gel composition |  |  |  |  |  |
| --- | --- | --- | --- | --- | --- | --- | --- | --- | --- | --- | --- |
| | Young's modulus (kPa) | % AAm (Con final) | % MBAA (Con final) | % APS (Con final) | % TEMED (V/V final) | total volume ($\mu\text{L}$) | volume $\text{dH}_2\text{O}$ ($\mu\text{L}$) | AAm 40% stock ($\mu\text{L}$) | MBAA 2% stock ($\mu\text{L}$) | APS 10% stock ($\mu\text{L}$) | TEMED 100% stock ($\mu\text{L}$) |
| 1 | $0.93 \pm 0.03$ | 4.00% | 0.17% | 0.25% | 0.25% | 10000 | 7875.00 | 1000.00 | 850.00 | 250.00 | 25.00 |
| 2 | $1.23 \pm 0.23$ | 4.94% | 0.10% | 0.25% | 0.25% | 10000 | 7999.80 | 1234.50 | 490.70 | 250.00 | 25.00 |
| 3 | $2.19 \pm 0.34$ | 5.00% | 0.15% | 0.25% | 0.25% | 10000 | 7737.05 | 1250.50 | 737.45 | 250.00 | 25.00 |
| 4 | $3.73 \pm 0.31$ | 5.50% | 0.23% | 0.25% | 0.25% | 10000 | 7200.00 | 1375.00 | 1150.00 | 250.00 | 25.00 |
| 5 | $0.90 \pm 0.11$ | 6.00% | 0.06% | 0.25% | 0.25% | 10000 | 7935.00 | 1500.00 | 290.00 | 250.00 | 25.00 |
| 6 | $7.45 \pm 0.28$ | 7.50% | 0.32% | 0.25% | 0.25% | 10000 | 6250.00 | 1875.00 | 1600.00 | 250.00 | 25.00 |
| 7 | $5.91 \pm 0.53$ | 9.00% | 0.14% | 0.25% | 0.25% | 10000 | 6775.00 | 2250.00 | 700.00 | 250.00 | 25.00 |
| 8 | $10.07 \pm 0.16$ | 10.00% | 0.20% | 0.25% | 0.25% | 10000 | 6225.00 | 2500.00 | 1000.00 | 250.00 | 25.00 |
| 9 | $14.46 \pm 0.06$ | 10.00% | 0.40% | 0.25% | 0.25% | 10000 | 5225.00 | 2500.00 | 2000.00 | 250.00 | 25.00 |
| 10 | $16.9 \pm 0.14$ | 10.00% | 0.43% | 0.25% | 0.25% | 10000 | 5075.00 | 2500.00 | 2150.00 | 250.00 | 25.00 |
| 11 | 20.94 ± 2.94 | 10.50% | 0.45% | 0.25% | 0.25% | 10000 | 4850.00 | 2625.00 | 2250.00 | 250.00 | 25.00 |
| 12 | 12.08 ± 1.95 | 12.00% | 0.20% | 0.25% | 0.25% | 10000 | 5725.00 | 3000.00 | 1000.00 | 250.00 | 25.00 |
| 13 | 23.56 ± 3.58 | 12.00% | 0.40% | 0.25% | 0.25% | 10000 | 4725.00 | 3000.00 | 2000.00 | 250.00 | 25.00 |
| 14 | 36.70 ± 4.30 | 18.00% | 0.40% | 0.25% | 0.25% | 10000 | 3225.00 | 4500.00 | 2000.00 | 250.00 | 25.00 |
| 15 | 47.25 ± 5.51 | 23.00% | 0.45% | 0.25% | 0.25% | 10000 | 1725.00 | 5750.00 | 2250.00 | 250.00 | 25.00 |
| 16 | 56.13 ± 6.82 | 28.90% | 0.50% | 0.25% | 0.25% | 10000 | 0.00 | 7225.00 | 2500.00 | 250.00 | 25.00 |

### Step-by-step method details

#### Synthesis of a large, unattached PAAm hydrogel

##### Timing: 1 h per gel + overnight swelling

In this step, a large circular unattached PAAm hydrogel with a flat surface is synthesized (Fig. 1 for schematic protocol, Fig. 2 for results). To this end, AAm monomer and MBAA crosslinker are mixed in varying concentrations, which determine the stiffness of the resulting PAAm gel. Radical polymerization is initiated by the addition of APS and TEMED, and the gel is cast between the lid and base dish of a circular polystyrene 92 mm Petri dish. These steps are performed on ice with precooled reagents to allow sufficient handling time. The polymerization itself is carried out at room temperature, as is customary in the field. The polymerized gel is then stamped into custom shapes and sizes and allowed to swell overnight.

**Figure 1:**
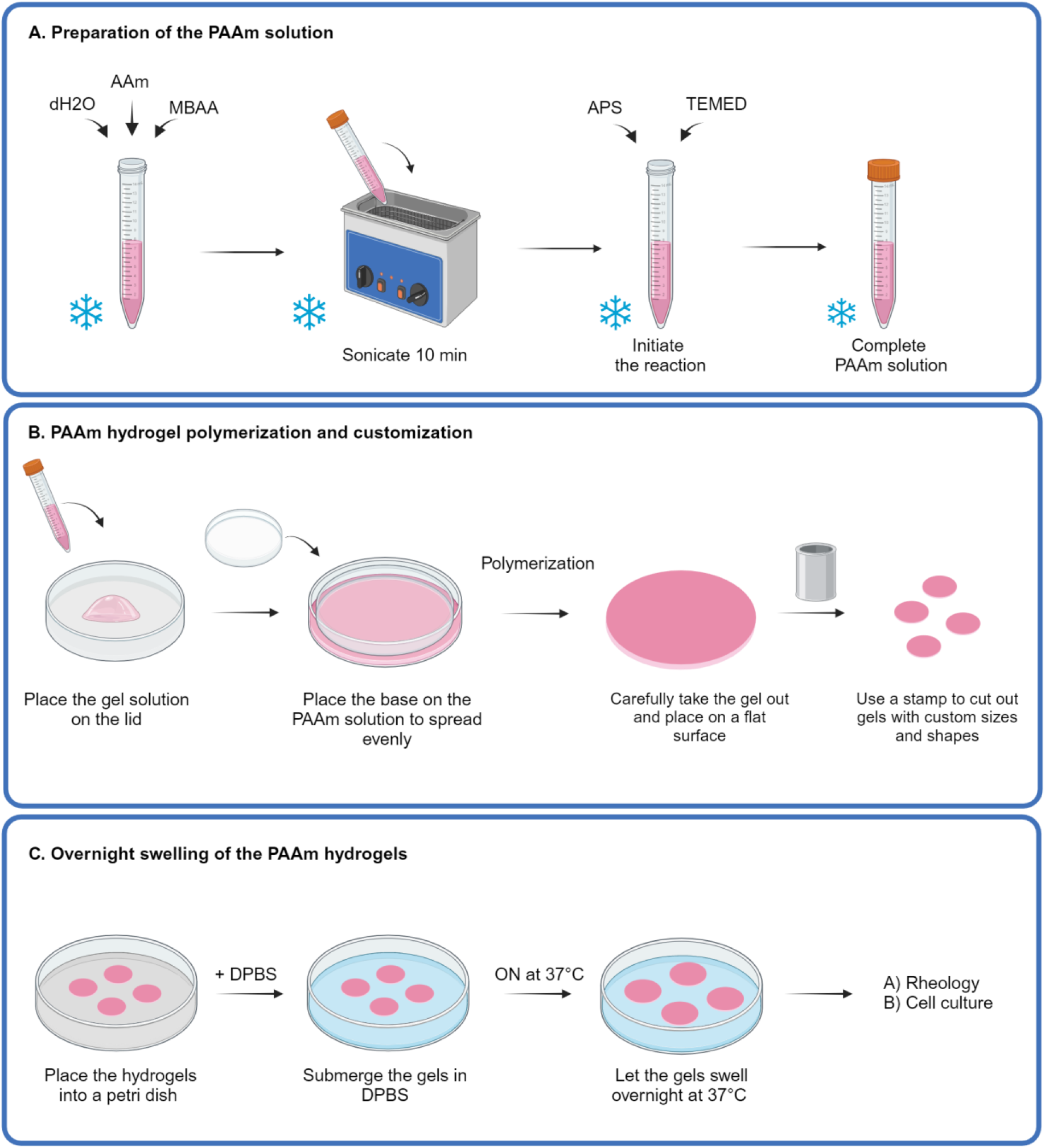
Production of a large, unattached PAAm hydrogel: scheme. **A:** Preparation of liquid PAAm mixture on ice by combination of distilled water (dH_2_O), AAm and MBAA, sonication, and addition of APS and TEMED. Cfr. steps 4-5 and 7. **B:** PAAm gel casting, polymerization, and customization. Cfr. steps 8-13. **C:** Overnight swelling. Cfr. step 16. Note that PAAm gels are normally transparent, but were artificially colored in this scheme for display purposes. Figure created with BioRender.com.

**Figure 2:**
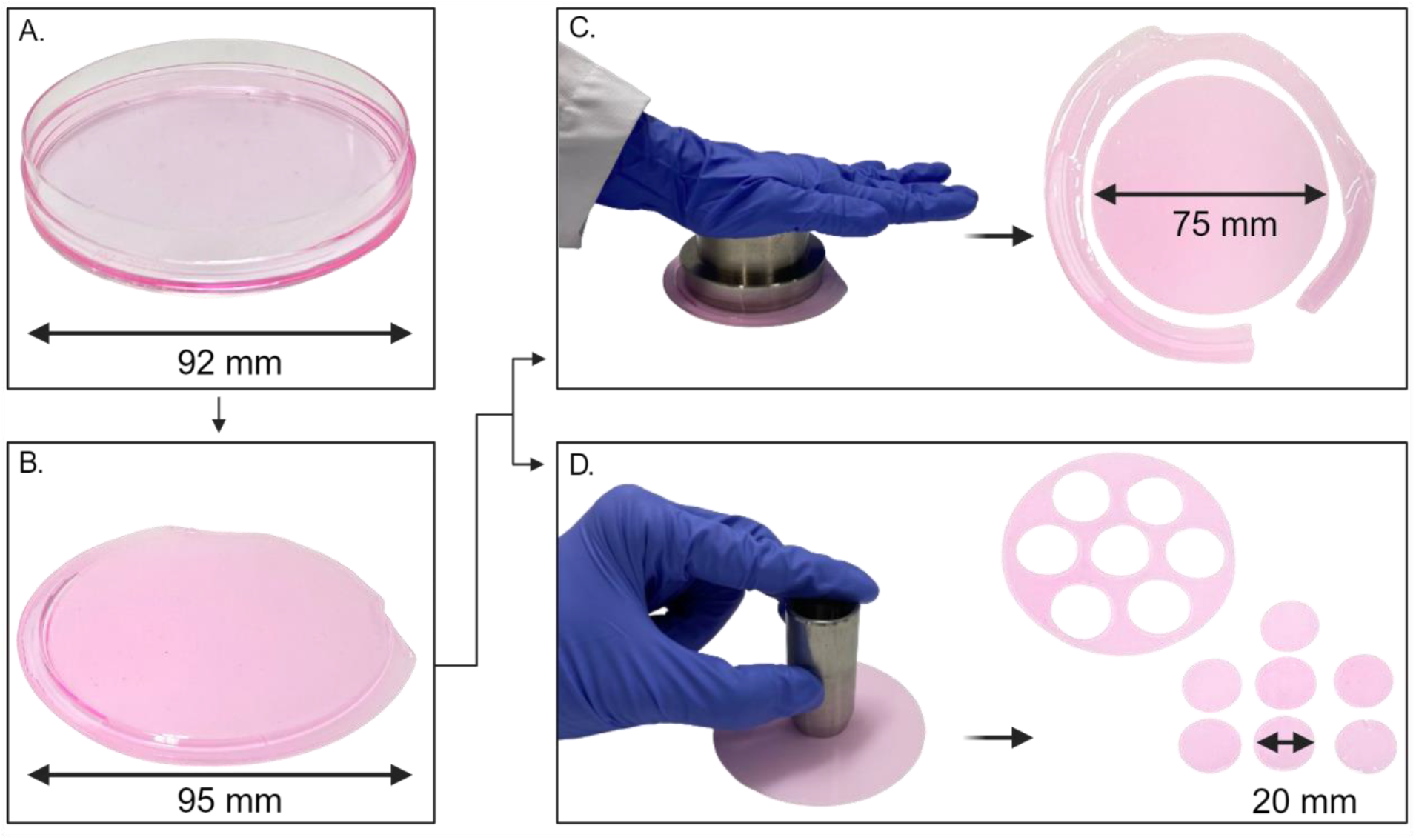
Production of a large, unattached PAAm hydrogel: practical results. **A:** PAAm gel cast in the polymerization chamber. Cfr. step 10. **B:** 95 mm unattached PAAm hydrogel before stamping. Cfr. step 12. **C**,**D:** Customized hydrogels after stamping, resulting in one 75 mm gel for large-batch cell culture applications (C), or 7 gel pieces of 20 mm for rheological characterization (D). Cfr. step 13. Note that PAAm gels are normally transparent, but were artificially colored in this image for display purposes. Figure created with BioRender.com.

**NOTE:** Work under a chemical fume hood, since several reagents are toxic upon ingestion/inhalation or represent a long-term health hazard.

**CRITICAL**: Since temperature affects the kinetics of PAAm polymerization, and therefore gel structure and stiffness, it is important to respect the indicated temperatures.^5^

**TIP:** It is possible to multiplex the production and customization of PAAm gels once one is familiar with the protocol. We recommend casting maximally 3 gels in parallel to avoid dehydration of polymerized gels. For example, it is possible to prepare a third PAAm mixture (cfr. step 4), and customize an already polymerized gel (cfr. steps 11 – 16c) while a second PAAm mixture is sonicating (cfr. step 5).

1. Pre-cool a circular polystyrene 92 mm Petri dish at 4 °C.
2. Place the vials containing dH_2_O, AAm 40% solution, MBAA 2% solution, APS 10% solution, TEMED 100% solution, and an empty 15 mL conical tube on ice.
3. Briefly vortex the AAm and MBAA solutions.
4. Add to the empty tube, on ice, the desired volumes of respectively dH_2_O, AAm and MBAA (Table 1, “practical gel composition”). Mix by gently pipetting up and down multiple times.
5. Sonicate the mixture on ice for 10 min. Longer sonication times are possible as long as the ice is still present, but we do not recommend sonicating longer than 20 min. **TIP**: Add water to the ice to improve the contact surface and cooling capacity. **CRITICAL**: After sonication, avoid vortexing, vigorous shaking or introducing air by pipetting, since oxygen acts as a free radical trap that inhibits PAAm polymerization.
6. Organize the polymerization chamber.
  a. Place the precooled 92 mm Petri dish at room temperature (23 °C).
  b. Place the lid with the open side up on a horizontal flat surface.
7. On ice, add the desired volumes of respectively APS and TEMED (Table 1, “practical gel composition”) to complete the PAAm reaction mixture. Mix gently by pipetting. **CRITICAL**: Polymerization is initiated once APS and TEMED are added, so it is imperative to work fast until the gel is cast. **NOTE**: As mentioned by Shi & Janmey, the final concentrations of APS (0.25% w/v) and TEMED (0.25% v/v) used here are higher than the commonly used standard.^1^ This is done to minimize mechanical perturbations that could arise before the polymerization is complete, when the base dish is floating without support on top of the liquid PAAm reaction mixture.
8. Transfer 9.5 mL of the complete PAAm liquid mixture to the open side of the Petri dish lid (Fig. 1B). **NOTE:** The transferred volume depends on the diameter of the Petri dish and may need to be optimized accordingly. A volume of 9.5 mL was found to be optimal for the 92 mm Petri dish used in this protocol. These experimental conditions yield gels with a diameter of approximately 95 mm, and a thickness of approximately 1.2 - 1.5 mm.
9. Close the polymerization chamber (Fig. 1B).
  a. Gently place the base dish with the open side up on top of the transferred solution. Start on one side and gently tilt until it touches the PAAm mixture everywhere and stands horizontally.
  b. Center the base dish with respect to the lid.
  c. Gently rotate the base dish approximately a quarter turn to ensure an equal distribution of the mixture throughout the polymerization chamber and its edges. **CRITICAL:** It is imperative to avoid air bubbles, as these will inhibit the polymerization and introduce heterogeneities in the gel.
10. Let the PAAm gel polymerize at room temperature (23 °C) (Figs. 1B and 2A). **CRITICAL:** Allow enough time for complete polymerization but avoid too long incubation periods to prevent dehydration. **TIP:** Place the remaining mixture in the tube at room temperature besides the polymerization chamber. By gently touching the mixture, one can determine when the reaction is complete. <u>Troubleshooting 1</u>. **NOTE:** Depending on the selected PAAm composition, polymerization could take between 5 to 20 min for gels listed in Table 1. Softer gels (≤ 20 kPa) may take significantly longer to polymerize than stiffer gels. In addition, polymerization may take longer when APS and TEMED are stored for prolonged periods of time (> 6 months) since this reduces their activity.
11. Once polymerization is finished, open the polymerization chamber.
  a. Carefully add 1 mL dH_2_O on the open edge of the base dish.
  b. Incubate a few minutes to allow the dH_2_O to spread.
  c. Gently pull on one side of the base dish to remove it.
  **NOTE:** The formed gel is not covalent attached to any surface and can readily be manipulated or transferred. Softer gels are more fragile and prone to rupture and need to be handled with great care. <u>Troubleshooting 2</u>. **NOTE:** In some instances, especially for softer gels (≤ 20 kPa), the PAAm gel can stick to the bottom side of the dish base, rather than remaining in the lid. In this case, place the Petri dish base with the gel oriented on the upper side on the bench and retrieve the gel from there.
12. Transfer the PAAm gel from the polymerization chamber onto a flat surface (Figs. 1B and 2B). **NOTE**: To prevent the stamp’s edge from wearing out, the gel is not stamped directly on the Petri dish lid. It is instead transferred to a surface made from a relatively soft material, for example a flat piece of polyvinyl chloride (PVC) plastic.
  a. Hold the lid upside down with the gel facing the flat surface.
  b. Gently push the round-tipped tweezer in between the Petri dish lid and the gel. Start from one edge and work towards the center of the gel, until the gel detaches from the lid and descends onto the flat surface. **TIP**: Hold the tweezer parallel to the lid surface as much as possible to avoid sharp contact with the gel and possible rupture.
  c. Spread out the gel evenly across the surface.
    i. Use the round-tipped tweezer and optional aid of scoop/spatula to remove wrinkles or folds.
    ii. Take special care of the edges, which tend to fold back onto the gel.
    **CRITICAL:** Visually check that the upper surface of the gel is not tilted. <u>Troubleshooting 3</u>.
13. Stamp the PAAm gel into one or multiple smaller pieces of custom shapes and sizes (Figs. 1B and 2C-D).
  a. Place a tissue-culture stamp or patisserie garnish stamp with sharp edge onto the gel.
  b. Press the stamp down with equal force from all sides using the palm of your hand. **NOTE**: Stiffer gels or larger stamps may require a considerable amount of force.
  c. Check that the stamp pushed through the gel on all sides by moving the tweezer around the stamp and seeing if it is unattached from the gel outside the stamp.
  **CRITICAL:** Avoid using the outer 0.5 cm of the large gel, since oxygen may have entered the polymerization chamber from the open edges. Oxygen is known to act as a free radical trap and inhibit the polymerization, possibly locally affecting the gels’ mechanical properties.^10^ **NOTE:** For large-batch cell culture applications (Major option B), the gel was stamped into one circular piece (diameter: 75 mm) (Fig. 2C). For rheological characterization (Major option A), the gel was stamped into 7 circular pieces (diameter: 20 mm) by making 1 stamp in the center and 6 stamps in a concentric ring around the center stamp (Fig. 2D).
14. Using the scoop/spatula and gentle sliding, transfer the gel piece(s) into a large container such as a large square or circular Petri dish.
15. Wash the piece(s) 3x with dH_2_O to remove unreacted PAAm monomers. The volume used should cover the whole gel and will depend on the size of the gel. For example, 2mL for a 20 mm circular gel, or 10 mL for a 75 mm circular gel.
16. Swell the gel(s) (Fig. 1C).
  a. Add DPBS 1x to fully submerge the gel(s). This is typically approximately 10x the volume used in step 15.
  b. Ensure the gel(s) is/are covered in liquid and not floating on top of it.
  c. Place the gel(s) in a humidified incubator at 37 °C for approximately 15 min.
  d. Ensure the gel(s) is/are still fully submerged and free-floating, otherwise readjust the position and/or add more DPBS 1x. This is necessary for gels that swell, since gels that are attached may wrinkle and rise above the liquid level when pushing themselves away from the Petri dish.
  e. Continue to swell the gel(s) overnight.

## Major option A: Rheological characterization

### Major step A1: Rheology measurements

#### Timing: 1 h per gel

In this step, a 20 mm swollen PAAm gel is mechanically characterized using a stress-controlled rotational rheometer with parallel plate geometry. Sanding paper is added to the rheometer components to enable grip on the already polymerized gel and prevent slipping during the rotational movements.

**CRITICAL:** Measurements are performed after overnight swelling to guarantee that the gels have reached the swelling equilibrium, a vital step as swelling leads to softening (decreased stiffness). For assessing stiffness of gels used in cell culture applications, where they remain immersed in cell culture medium or buffers for extended periods, pre-swelling the gels in DPBS at 37 °C is essential.

**CRITICAL:** Measurements can be performed on gels of various sizes as long as they fit the rheometer’s dimensions, and the reagents are proportionally adjusted. It is important to remember that using sizes other than the standard 20 mm may lead to small variations in stiffness values due to edge effects at the periphery of the geometry.

**TIP:** Multiple gels can be characterized sequentially, provided that the sandpapers are replaced whenever grip issues emerge, or at a minimum, at the start of every new day.

17. Switch on rheometer hardware, including the water pump. **NOTE:** Ensure the water reservoir is filled. In case the error ‘low amount of water’ is visible, refill with dH_2_O.
18. Switch on software Trios version 5.5.0.323.
19. Attach sandpaper to the geometry and lower plate.
  a. Customize sandpaper (Fig. 3A).

**Figure 3:**
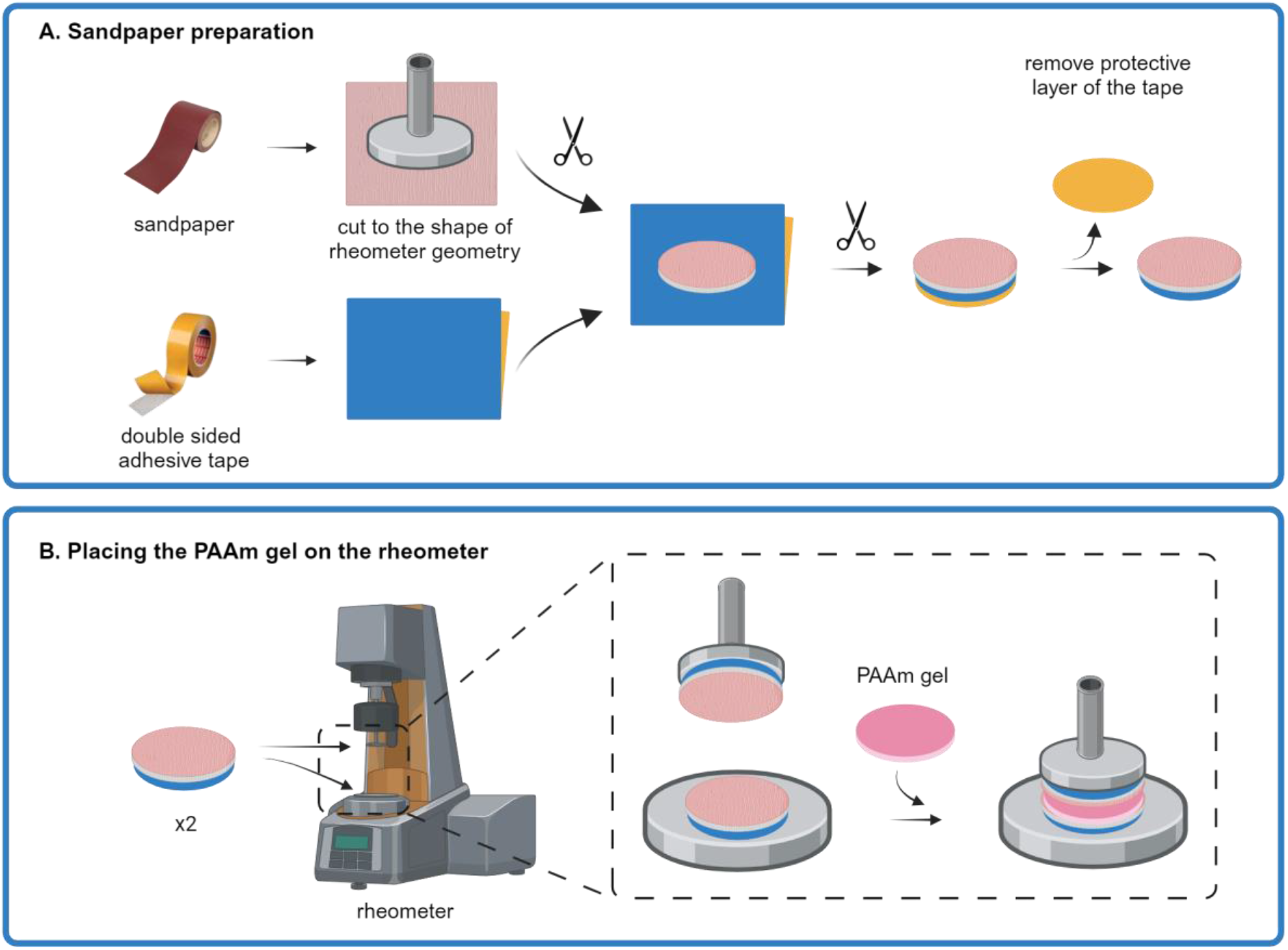
Attachment of sandpaper on rheometer components. **A:** Sandpaper customization. Cfr. step 19a. **B:** Attach customized sandpaper to double-sided adhesive tape Cfr. step 19b. **C-D:** Attach sandpaper to geometry and lower plate, and place the gel on the rheometer. Cfr. steps 19d, 19f, and 22i. Note that PAAm gels are normally transparent, but were artificially colored in this scheme for display purposes. Figure created with BioRender.com.
    i. Draw a circle with 20 mm diameter on the back of the sandpaper using a pencil. **TIP:** Outline the shape of the 20 mm geometry by placing it on the back of the sandpaper.
    ii. Cut out the circle as precisely as possible using scissors. Use micro-scissors for the best result.
  b. Attach customized sandpaper to double-sided adhesive tape (Fig. 3A).
    i. Cut a piece of double-sided tape with dimensions larger than a 20 mm circle.
    ii. Stick the sandpaper circle to the exposed adhesive side of the tape.
    iii. Cut the tape to the same size as the sandpaper as precisely as possible.
  c. Repeat steps a and b to create a second circle of sandpaper attached to double-sided tape.
  d. Attach the first sandpaper circle to the geometry (Fig. 3B).
    i. Remove the protective layer from the double-sided tape.
    ii. Lay on a flat surface with the sandpaper facing downwards and the adhesive side upwards.
    iii. Place the geometry as accurately as possible on top.
    iv. Push the geometry down firmly with equally distributed force.
    v. Smoothen the edges by cutting away protrusions of sandpaper and/or adhesive tape. **CRITICAL:** Meticulously remove remaining tape and sandpaper from the edges of the geometry, since any disturbance of the geometries’ outer edges will lead to aberrant results.
  e. Install the geometry on the rheometer.
    i. Press the “lock” button until the rheometer beeps.
    ii. Slide the geometry up the driving shaft.
    iii. Align geometry calibration line with the calibration line on the rheometer.
    iv. Hold the geometry stationary and turn the draw rod clockwise to secure it.
    v. Press the “lock” button.
    vi. Observe the geometry rotating freely.
    vii. In the software, check that the correct geometry serial number is selected.
  f. Attach the second sandpaper circle to the lower plate (Fig. 3B).
    i. Lower the geometry using arrows on the keypad until it almost touches the lower plate.
    ii. With a pencil, draw the outline of the geometry on the lower plate.
    iii. Move the geometry back to the upper position.
    iv. Remove the protective layer from the double-sided tape.
    v. Paste it as accurately as possible in the drawn outline on the lower plate.
    vi. Press firmly with equally distributed force.
  g. Visually check that both sandpapers are positioned correctly. <u>Troubleshooting 4</u>.
    i. Lower the geometry using the arrows on the keypad.
    ii. The two pieces of sandpaper on the geometry and lower plate should align exactly. **CRITICAL:** Customization and attachment of the sandpaper is extremely important since any disturbance of the geometries’ outer edges or lower plate will lead to aberrant results.
  h. Hydrate sandpapers.
    i. Raise the geometry slightly.
    ii. Incubate 1 mL DPBS 1x between both sandpapers for 5 min.
    iii. Gently remove the excess with a precision wipe. **NOTE:** This step is necessary to prevent the sandpaper from dehydrating the PAAm gel.
20. Calibrate the rheometer.
  a. Set rheometer temperature to 21 °C. **NOTE:** This temperature differs from the 37 °C used in cell culture. However, increasing the experiment above the room temperature would require use of a humidity chamber that prevents the user from observing the gel while performing the measurement. Since gel positioning and observation of contact between the gel and geometry is critical for this experiment, and since bulk mechanical properties measured on a rheometer were previously reported to be identical when measured at 21 °C and at 37 °C, we decided to prioritize this requirement above the incubation temperature.^6^
  b. Perform inertia calibration in the software.
  c. Perform friction calibration in the software.
  d. Perform rotational mapping calibration with bearing mapping type “standard” and number of iterations “2” selected.
  e. Perform zero-gap calibration. **CRITICAL:** The geometry will automatically be lowered, so keep the rheometer area clear. **NOTE for steps b-e:** Calibrations are performed only after attachment and hydration of the sandpaper, since the altered weight (distribution) and sandpaper thickness will affect the calibration.
21. Set up rheology measurement protocol in the software according to parameters in Table 2.

**Table 2:** Rheology acquisition protocol.
  a. Parameters to set for the different measurement steps include:
    i. “Environment”: temperature.
    ii. “Soak time”: incubation time before starting the measurement, important when the gel needs to relax in between intensive measurements.
    iii. “Duration”: duration of the measurement in case of a time sweep.
    iv. “Strain”: amplitude or displacement of the rotational movements, reflecting how large the angle of the geometry rotations is.
    v. “Frequency”: how fast the geometry is rotated.
    vi. “Measurement type”: if and how duration, strain or frequency are altered during the measurement.
  b. This protocol will perform 4 different measurement steps, namely:
    i. 30 sec time sweep at 1% strain, 1 Hz frequency.
    ii. 30 sec time sweep at 2% strain, 1 Hz frequency.
    iii. Logarithmic frequency sweep from 0.016 Hz to 1.592 Hz frequency, at 1% strain.
    iv. Logarithmic amplitude sweep from 0.01% to 10% strain, at 1.0 Hz frequency. **NOTE:** As the larger deformations in the strain sweep step may damage the gel, it is recommended to add it as the final step of the protocol. The order of the other steps should not influence its outcomes.
22. Perform rheology measurement.
  a. Transfer a 20 mm PAAm gel that was swollen overnight onto a flat PVC surface.
  b. Restamp the gel to a 20 mm circle, as its dimensions could have been altered by gel swelling. **CRITICAL:** This restamping is essential to ensure matching of gel and geometry dimensions.
  c. Ensure sandpapers are still hydrated but not soaked.
  d. With geometry in the upper position and able to rotate freely, calibrate axial force. Axial force should be an order of magnitude 10^−3^ or 10^−4^ N after calibration.
  e. Place the gel in the center of the sandpaper on the lower plate (Fig. 3B).
  f. Remove the excess DPBS on the sides with a precision wipe.
  g. Lower the geometry using the keypad till just above the gel (gap 2000 nm).
  h. Verify that the gel is placed centrally with respect to the geometry. Adjust its position by careful sliding if necessary.
  i. Lower the geometry in steps of 10-50 nm in the software until it touches the PAAm gel everywhere (Fig. 3B). **CRITICAL:** It is extremely important that the geometry touches the PAAm gel on all sides (left, right, front, back), check this visually. Failure to do this will result in uneven geometry rotation throughout the sample and hence aberrant results.
  j. Lower the geometry in steps of 10 nm until the axial force is no longer negligible (i.e. minimally 0.01 N). Write down the initial gap distance and initial axial force value for this “initial gap”. **NOTE:** Axial force may already be higher than 0.2 N because of the requirement of uniform gel contact in step 22i. This may indicate that the sandpaper or gel surface is tilted and should be avoided.
  k. Compress the gel 3% by lowering the gap to the “final gap”. Calculate the final gap distance by the following formula. Take note of the final gap distance and corresponding final axial force value.

| Measurement step | Measurement name | Environment | Soak time | Duration | Strain | Frequency | Measurement type |
| --- | --- | --- | --- | --- | --- | --- | --- |
| start | Conditioning options: disabled | NA | NA | NA | NA | NA | NA |
| 1 | Time sweep at 1% strain | 21 °C | 0" | 30" | 1% | 1.0 Hz<br>= 6.283 rad/s | Single point |
| 2 | Time sweep at 2% strain | 21 °C | 0" | 30" | 2% | 1.0 Hz<br>= 6.283 rad/s | Single point |
| 3 | Frequency sweep | 21 °C | 30" |  | 1% | 0.0159155 Hz to<br>1.59155 Hz<br>= 0.1 rad/s to 10.0 rad/s | Log sweep |
| 4 | Amplitude sweep | 21 °C | 30" |  | 0.01% to<br>10% | 1.0 Hz<br>= 6.283 rad/s | Log sweep |
| end |  | Set temperature idle | NA | NA | NA | NA | NA |

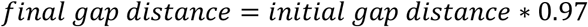

**NOTE:** This compression step is necessary to ensure that the geometry and lower plate indeed contact the gel, and do not measure a thin water film between the gel and the geometry, which would result in an aberrant stiffness measurement.
  l. Zero the axial force. Axial force should then be an order of magnitude 10^−3^ or 10^−4^ N.
  m. Start the measurement. **CRITICAL:** In the point displacement curve, note that its amplitude and period match that of the torque curve, but with a slight time delay. This indicates proper gripping of the geometry on the PAAm gel.
  n. Upon completion of the measurement, raise the geometry to the upper position and remove the gel.
23. Repeat measurements on a second and third 20 mm technical replicate from the same gel.
24. Export the data as an excel file.
  a. The measurement steps are saved in different tabs of the excel file called “Time sweep –1”, “Time sweep – 2”, “Frequency sweep – 3” and “Amplitude sweep – 4”.
  b. Rheometer specifications are saved in the tab “Details”.

## Major step A2: Rheology data analysis

### Timing: 20 min per gel

Rheology data is analyzed to obtain information about the frequency- and amplitude-dependency of the gels. Additionally, the gel’s stiffness (Young’s modulus) is derived from the storage modulus, allowing us to describe this property in function of the AAm and MBAA content.

25. Time sweeps and stiffness.
  a. Visualize results of times sweeps at 1% and 2% strain. Plot storage modulus (*G*′, in Pa) and loss modulus (*G*′′, in Pa) in function of time (t, in seconds) using a 2D line graph with markers and y-axis in logscale (Fig. 4A for result).

**Figure 4:**
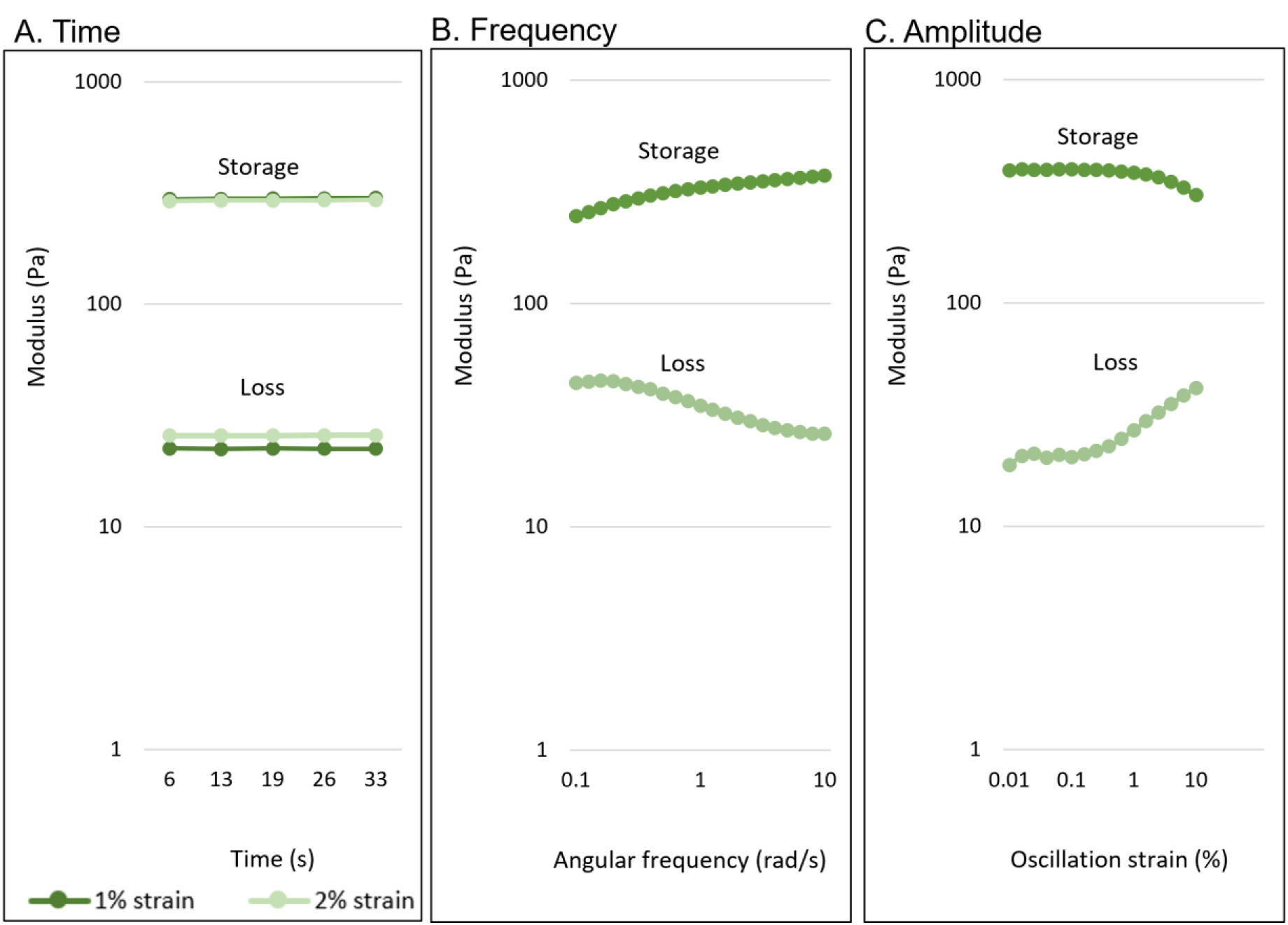
Examples of PAAm rheology results. Storage and loss moduli measured during time sweeps at 1% and 2% strain (**A**) Cfr. step 25a, frequency sweeps (**B**) Cfr. step 26a and amplitude sweeps (**C**) Cfr. step 27a are shown.

**Figure 5:**
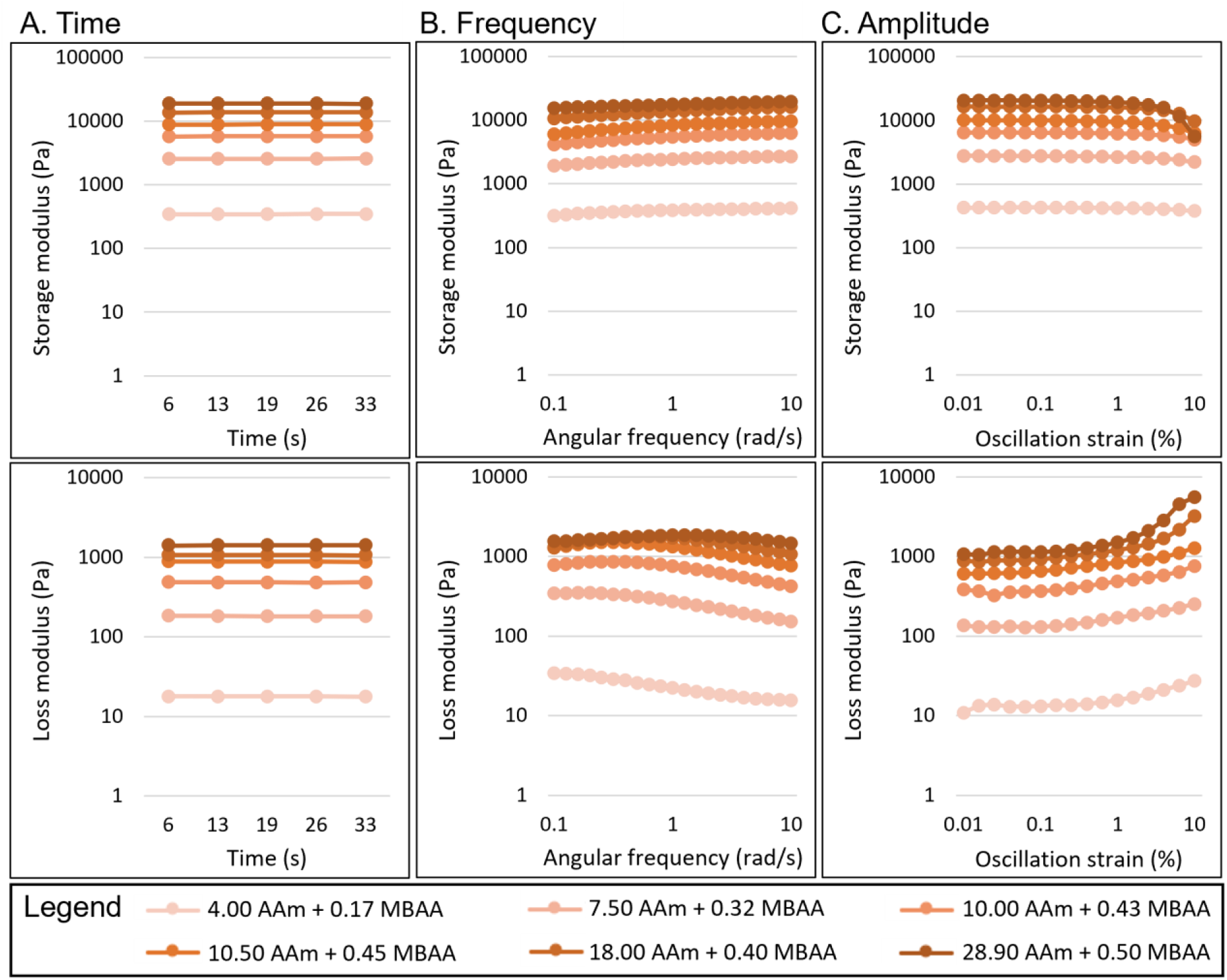
Rheological characterization of PAAm library. Storage and loss moduli measured during time sweeps at 1% strain (A), frequency sweeps (B) and amplitude sweeps (C) are shown for some of the characterized gel compositions. Note that scale bars differ between storage and loss modulus. Cfr. section “Expected outcomes”.
  b. Calculate the average stiffness of the PAAm gel from at least 3 technical replicates from the same large gel.
    i. Per measurement, calculate the average storage modulus (*G*′, in Pa) across the different timepoints for 1% strain.
    ii. Per measurement, calculate the stiffness (Young’s modulus, E, in kPa):

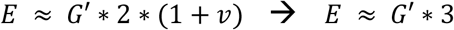
    iii. Where v is Poisson’s ratio which describes the compressibility of a material and is assumed to be 0.5 for PAAm gels.^3,4,11^
    iv. Calculate the average gel stiffness (E_average_) from the technical replicates.
    v. Calculate the percentage error on the gel stiffness:

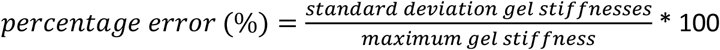
    vi. Calculate the absolute error on the gel stiffness:

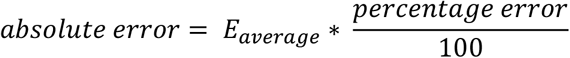
26. Frequency sweep.
  a. Visualize results of the frequency sweep. Plot storage modulus (*G*′, in Pa) and loss modulus (*G*′′, in Pa) in function of angular frequency (ω, in rad/sec or Hz) using a 2D line graph with markers and y-axis in logscale (Fig. 4B for result). **NOTE:** rad/sec and Hz can be converted since 1.0 Hz ≈ 6.283 rad/sec.
27. Amplitude sweep.
  a. Visualize results of the amplitude sweep. Plot storage modulus (*G*′, in Pa) and loss modulus (*G*′′, in Pa) in function of oscillation strain (γ, in %) using a 2D line graph with markers and y-axis in logscale (Fig. 4C for result).

## Major option B: Cell culture applications

### Major step B1: Hydrogel functionalization

#### Timing: 45 min per gel + overnight incubation + 45 min per gel

To enable cell attachment on biologically inert PAAm gels, their surface is functionalized with extracellular matrix (ECM) molecules. While several alternatives have been described, we use the bi-functional crosslinker sulfo-SANPAH in a photo-activated reaction using UV-light to covalently link collagen type I to the gel surface.^6,12^

**NOTE**: During the UV-treatment for sulfo-SANPAH functionalization gels are sterilized, and all subsequent steps should be done in a sterile environment (step 33 and further).

**CRITICAL:** Always verify that the gel is submerged and not floating on top of added solutions.

**CRITICAL:** Quantities are specified as product amount (µg) per surface area (cm^2^) since products react with the surface, making liquid volumes (mL) less critical as long as surfaces are fully covered. The protocol details procedures for a 75 mm circular gel, used in large-batch cell culture, but can be adapted for different shapes and sizes when µg/cm^2^ ratios are maintained.

**TIP:** Once familiar with the protocol, it is feasible to multiplex functionalization. During the first sulfo-SANPAH incubation (step 33), a previously functionalized gel can undergo collagen incubation (steps 35-37b); and new gel can be stamped and washed (steps 28-32) during the second sulfo-SANPAH step (step 34). However, avoid extending sulfo-SANPAH incubation times and prevent hydrogel dehydration.

28. Remove DPBS 1x from the container.
29. Transfer the PAAm gel onto a flat PVC surface.
30. Restamp the gel to a 75 mm circle, as its dimensions are affected by gel swelling. **NOTE:** Gels of varying stiffnesses swell differently depending on their stiffness, up to 130% in diameter for the gels produced here.
31. Transfer the gel to a 92 × 16 mm circular PS Petri dish.
32. Wash the gel 3x with 10 mL dH_2_O.
33. Incubate with 0.1 mg/cm^2^ sulfo-SANPAH.
  a. Mix 6.66 µL sulfo-SANPAH 100 mg/mL (prepared using Recipe 4) with 10 mL sterile dH_2_O.
  b. Incubate on the gel.
  c. Ensure the liquid is spread homogenously over the entire Petri dish area and the gel is immersed entirely.
  d. Incubate at room temperature under 365 nm UV-light for 15 min, alongside the empty Petri dish lid for sterilization.
  e. Remove the sulfo-SANPAH solution. Gel surface should appear rust brown. **NOTE:** Sulfo-SANPAH is light and moisture sensitive. Protect stock solution from light and work in the dark. Prepare work solution on the spot and incubate immediately on the gel. **NOTE:** Sulfo-SANPAH concentration may influence the density of ECM ligands bound during the functionalization and may need to be optimized depending on the type of ECM ligand and the cell line employed in downstream applications.
34. Repeat previous step for a second round of functionalization.
35. Wash the gel 3x with 10 mL dH_2_O to remove unreacted sulfo-SANPAH. **NOTE:** Sulfo-SANPAH aggregates, visible as red thread-like structures, may be present. Attempt to remove using the pipet.
36. Transfer the gel to a new 92 × 16 mm circular PS Petri dish.
37. Incubate with 25 µg/cm^2^ collagen type I.
  a. Mix 55 µL collagen type I 3 mg/mL with 10 mL sterile 0.5% acetic acid solution with pH 3.40 (prepared using Recipe 6).
  b. Pipet on the gel.
  c. Incubate overnight on a rocker plate with seesaw movement at approximately 10 movements/minute, in the dark at room temperature. **NOTE:** Collagen type I is derived from animal origin and the concentration of purchased stock solutions may therefore deviate. Prepare work solution so that 165 µg collagen is present for a Petri dish with 92 mm diameter. **NOTE:** Collagen type I is diluted in 0.5% acetic acid at acidic pH to improve collagen heterogeneity and prevent aggregate formation. ^13^
38. Remove the collagen solution. **NOTE:** This solution may appear slightly red due to the excess sulfo-SANPAH that detached. The gel itself should look almost transparent.
39. Wash the gel 3x with 10 mL sterile DPBS 1x to remove unbound collagen type I.
40. Equilibrate the gel.
  a. Incubate the gel in 10 mL sterile prewarmed cell culture medium.
  b. Place in a humified incubator at 37 °C and 5% CO_2_ for 30 min. **TIP:** If needed, incubation can be extended up to 4 h.

### Major step B2: Cell seeding and growing

#### Timing: 45 min per gel + variable growth period (e.g. overnight)

KM12L4a cells are seeded on top of the functionalized PAAm gel and grown in this 2.5D setup for the desired time period, depending on the downstream application.

**NOTE:** Cell densities and incubation times may need to be adjusted depending on the applied cell type. <u>Troubleshooting 5</u>.

41. Prewarm the heating plate to 37 °C.
42. Harvest KM12L4a cells grown to 80% confluency according to standard cell culture practices using 1X trypsin-EDTA (prepared using Recipe 8).
43. Count cells according to standard cell culture practices using trypan blue and cell counting chamber.
44. Seed 10 million KM12L4a cells on a 75 mm functionalized gel.
  a. Dilute the cell suspension to 1 million cells per mL in cell culture medium (prepared using Recipe 7).
  b. Remove medium from the equilibrated gel and place it on the 37 °C heating plate.
  c. Place the stamp around the gel.
  d. Slowly pipet 10 mL cell suspension on top of the PAAm gel. **NOTE:** While some fluid may leak from the stamp, it will largely prevent cells from flowing off the gel.
  e. Gently move the Petri dish from right to left, and front to back to distribute cells homogenously.
  f. Incubate the gel to allow cell settling and initial attachment, check attachment and density on a microscope with transmission light (Fig. 6A).

**Figure 6:**
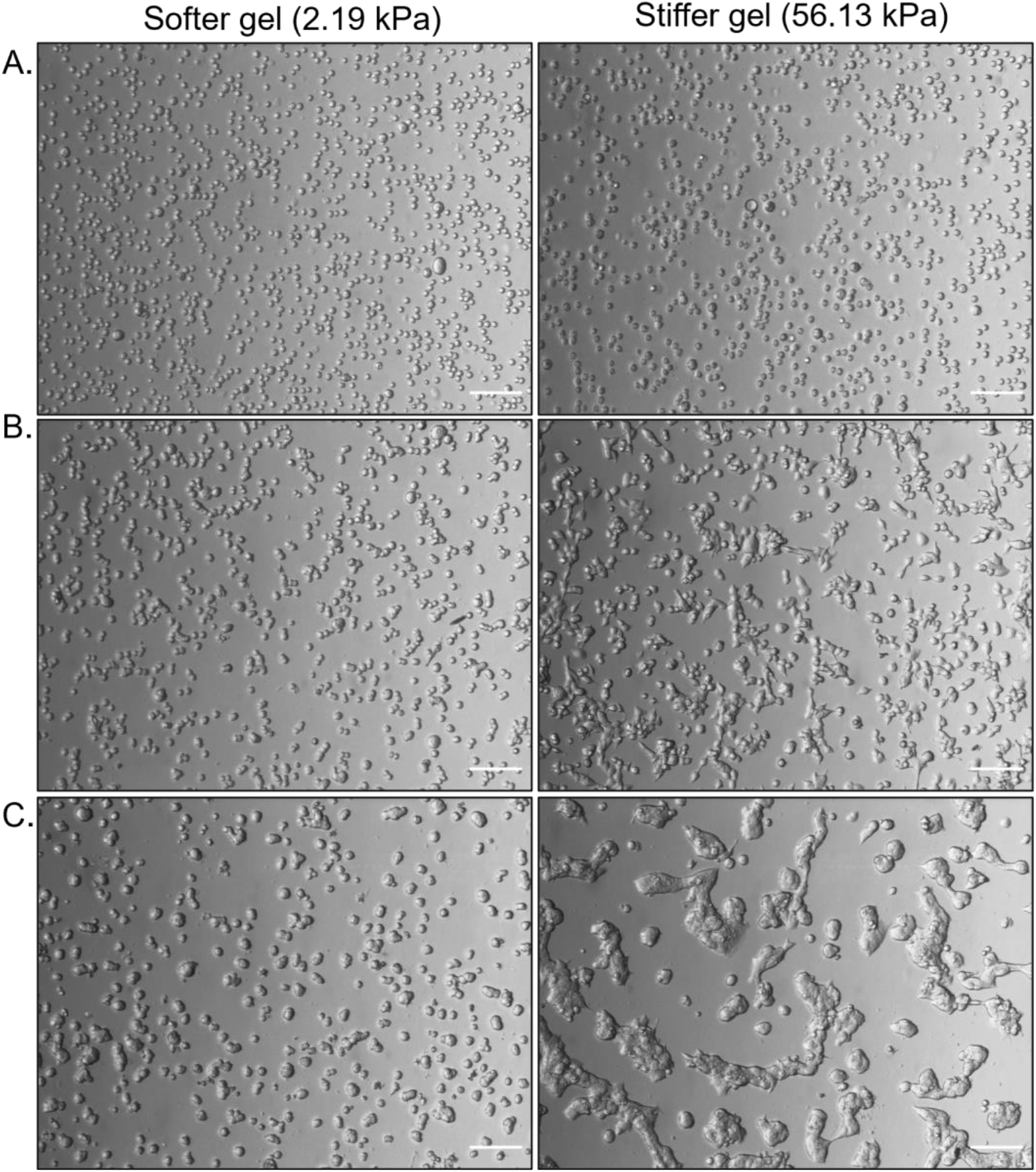
Growth of KM12L4a cells on PAAm gels of different stiffnesses on different timepoints. Images show cells just after cell seeding (A) Cfr. step 44f, after 24 h cell growth (B) Cfr. step 46, and after 48 h starvation (C) Cfr. step 51f. Images are from different regions of the same 75 mm PAAm gel. Scale bars 200 µm. **NOTE:** Incubation for 20 min is usually sufficient for stiffer gels (> 20 kPa), while 30 min are recommended for softer gels (≤ 20 kPa). <u>Troubleshooting 5</u>.
45. Grow cells for 24 h in a humified incubator at 37 °C and 5% CO_2_.

**NOTE:** Cells can be grown for variable time intervals depending on the downstream application.

## Downstream application 1: Secretome and cell collection

### Timing for starvation: 15 min per gel + 1 h incubation + 10 min per gel + 48 h incubation. Timing for secretome and cell harvesting: 45 min per gel

The secretome (conditioned medium) and cells can be harvested from the 75 mm gel and can be used for several applications including purification and analysis of secretome/exosomes, omics analyses, or *in vivo* experiments.

**CRITICAL:** Always verify that the gel is submerged and not floating on top of added solutions.

**NOTE:** When harvesting cells or secretome for proteomics applications, use commercial DBPS 1x to obtain repeatable conditions and to avoid the presence of potential protein contaminants, rather than the DPBS prepared using Recipe 1.

46. Check cell attachment and morphology on a microscope with transmission light (Fig. 6B). **NOTE:** Dead, floating cells can be present but should not dominate. They will be washed away before incubation with medium.
47. Remove cell culture medium.
48. Wash 2x with 10 mL prewarmed commercial DPBS 1x.
49. Transfer the gel to a new 92 mm Petri dish. **NOTE:** The gel is moved to eliminate cells that had flowed off the gel and grow on the plastic base of the Petri dish. This step is crucial when studying the impact of substrate stiffness since mixing cells grown on plastic with those grown on the hydrogel will introduce inaccuracies and could obscure differences between conditions.
50. Wash 2x with 10 mL prewarmed commercial DPBS 1x.
51. *Optional:* Starve cells for 48 h in serum-free medium. **NOTE:** Starvation is performed when collecting secretome (step 52), or cells for certain downstream analyses (step 53) when avoiding interference of FBS-components is critical. Depending on the application, this step may be omitted when harvesting cells for other purposes.
  a. Add 13 mL prewarmed starvation medium (prepared using Recipe 7).
  b. Incubate 1 h in a humified incubator at 37 °C and 5% CO_2_.
  c. Wash 3x with 10 mL prewarmed commercial DPBS 1x.
  d. Add 13 mL prewarmed starvation medium.
  e. Incubate 48 h in a humified incubator at 37 °C and 5% CO_2_.
  f. Check cell attachment and morphology on a microscope with transmission light (Fig. 6C). **NOTE:** KM12L4a cell morphology is not affected by 48 h starvation in our experience.
52. *Optional:* Harvest secretome.
  a. Collect the medium (secretome) in a 15 mL conical tube.
  b. Centrifuge 5 min at 140 g to pellet remaining cells and cell debris.
  c. Transfer the supernatant to a new 15 mL conical tube without disturbing the pellet.
  d. Freeze the secretome at -80 °C for further analysis.
53. Harvest cells.
  a. Place the Petri dish containing the gel on a 37 °C heating plate.
  b. Wash 3x with 10 mL prewarmed commercial DPBS 1x.
  c. Add 10 mL prewarmed trypsin-EDTA 1x solution (prepared using Recipe 8). **CRITICAL:** For proteomics, detach cells using sterilized 4 mM EDTA in commercial DPBS 1x.
  d. Incubate 2-3 min in a humified incubator at 37 °C and 5% CO_2_.
  e. Flow the solution over the gel multiple times to facilitate cell detachment. **CRITICAL:** Do not incubate trypsin-EDTA for longer than 5 min, and EDTA for longer than 30 min. **TIP:** Use a 1 mL micropipette to gently flow the solution in vertical lines across the gel surface. Rotate the Petri dish 90 ° and repeat to cover the whole surface. Be cautious to avoid liquid splashing, especially with softer gels (≤ 20 kPa).
  f. Confirm detachment of the majority of cells using transmission light on a microscope.
  g. Collect the cell suspension in a 15 mL conical tube.
  h. Retrieve the remaining cells by flowing an additional 3 mL prewarmed commercial DPBS 1x across the surface and collect in the same conical tube.
  i. Centrifuge 5 min at 140 g to pellet cells.
  j. Carefully remove supernatant without disturbing the cell pellet.
  k. Resuspend cells in 500 µL commercial DBPS 1x, simultaneously rinsing the tube wall.
  l. Transfer the cell suspension to a 1.5 mL Eppendorf tube. Measure the suspension volume (mL) by fully aspirating the liquid with the pipette.
  m. Count cells according to standard cell culture practices using trypan blue and cell counting chamber.
  n. Calculate the total number of harvested cells using the suspension volume measured in step 53l.
  o. Centrifuge 5 min at 140 g to pellet cells.
  p. Carefully remove supernatant without disturbing the cell pellet.
  q. Freeze the cell pellet at -80 °C for further analysis.

## Downstream application 2: Protein visualization by immunolabeling and fluorescence microscopy

### Timing for immunolabeling: 90 min + 2 h incubation + 10 min + overnight incubation + 20 min + 2 h incubation + 1 h

#### Timing for microscopy: 1 h

Instead of secretome and cell harvesting (Downstream application 1), cells grown on top of PAAm gels can be immunolabeled and visualized using confocal fluorescence microscopy to analyze the expression and subcellular localization of specific proteins. A protocol for immunolabeling of organoids embedded in Matrigel is adopted to ensure effective antibody penetration into densely grown KM12L4a cell clumps.^14^ Here, we immunolabel cytoskeletal alfa tubulin using a primary and secondary antibody, and visualize the nuclei with DAPI. When using other cell types or targets, the permeabilization time, antibody concentrations, incubation times and temperatures may need to be adjusted.

**CRITICAL:** Always verify that the gel is submerged and not floating on top of added solutions.

**CRITICAL:** Maintain temperatures of buffers (prewarmed to 37 °C) and incubation steps as indicated in the protocol to ensure efficient antibody penetration.

**CRITICAL:** Secondary antibodies and DAPI are light-sensitive, so protect from light upon storage and handling.

**NOTE:** In this protocol 75 mm gels are fixed, then restamped into 10 mm circular gels and placed in a 12 mm chamber with a recess of 4 mm, which allows the insertion of the round-tipped tweezers. We have opted to use these small gels to limit the use of reagents. Variously sized gels and chambers can be used provided that the reagents are adjusted proportionally, and the gel is fully submerged in the solutions. Different immunolabeling conditions can be tested in parallel on different 10 mm samples, and the remainder of the gel can be stored submerged in DPBS 1x at 4 °C for up to 2 weeks for subsequent tests (same biological replicate).

54. Remove cell culture medium. **NOTE:** Cells can be immunolabeled either after 24 h of growth or after a customized incubation time (with or without starvation medium), depending on the research question. Reagent quantities below are optimized for immunolabeling of KM12L4a cells after 24 h growth and 48 h starvation.
55. Wash 3x with 10 mL prewarmed DPBS 1x.
56. Fix the sample in 10 mL prewarmed PFA 4% (prepared using Recipe 9) for 1 h in a humified incubator at 37 °C and 5% CO_2_.
57. Remove PFA and wash 3x with 10 mL prewarmed DPBS 1x. **TIP:** The fixed sample can be stored in DPBS at 4 °C for up to 2 weeks.
58. Transfer the gel onto a flat PVC surface.
59. Restamp the gel to a 10 mm circle.
60. Transfer the gel to the immunolabeling chamber.
61. Permeabilize in 200 µL prewarmed Triton X-100 0.1% (prepared using Recipe 10) for 30 min at 4 °C. **NOTE:** Permeabilization is only required for immunolabeling of targets located intracellularly and could be omitted for extracellular antibody epitopes or when labeling with cell-permeable small molecules.
62. Remove Triton X-100 and wash 3x with 200 µL prewarmed DPBS 1x.
63. Block in 200 µL prewarmed blocking buffer (prepared using Recipe 11) for 2 h in a humified incubator at 37 °C and 5% CO_2_. **TIP:** If needed, incubation can be extended up to 4 h.
64. Remove blocking buffer.
65. Dilute 1 µg anti-tubulin primary antibody in 200 µL prewarmed blocking buffer.
66. Incubate primary antibody on the sample, overnight at room temperature.
67. Remove primary antibody and wash 3x with 200 µL prewarmed DPBS 1x. **TIP:** Primary antibody can be recycled up to 3 times when stored at 4 °C.
68. While protected from light, dilute 0.2 µg goat-anti-mouse Atto647N secondary antibody in 200 µL prewarmed blocking buffer. **NOTE:** The red-emitting Atto647N dye was selected because the longer excitation wavelength penetrates better through the cell clumps.
69. Incubate secondary antibody on the sample, 2 h at room temperature protected from light.
70. Remove secondary antibody and wash 3x with 200 µL prewarmed DPBS 1x for 5 min each.
71. Dilute 1 µg DAPI nuclear stain (take 0.2 µL of the 5 mg/mL stock solution prepared using Recipe 12) in 200 µL prewarmed blocking buffer.
72. Incubate DAPI on the sample, 15 min protected from light in a humified incubator at 37 °C and 5% CO_2_.
73. Remove DAPI and wash 3x with 200 µL prewarmed DPBS 1x for 10 min each in a humified incubator at 37 °C and 5% CO_2_. **TIP:** This sample can be stored at 4 °C for up to 4 days.
74. Place the sample inverted in a 6-well glass-bottom plate for microscopy imaging. **TIP:** Use the round-tipped tweezer and scoop/spatula to flip the gel upside down in the plate. **NOTE:** The sample is inverted so that cells are positioned closest to the coverslip, with the gel on top. This enables imaging of the cells with high resolution objectives that have a limited working distance and cannot reach the cells on top of the gel sample.
75. Add approximately 1.2 mL DPBS 1x so that the sample is surrounded by liquid, but not floating inside the well. <u>Troubleshooting 6</u>.
76. Startup Leica TCS SP8 confocal microscope hardware and software.
77. Place a drop of Milli-Q water on the 63x water objective.
78. Place the 6-well glass-bottom plate in the plate holder and onto the microscope stage.
79. Adjust image acquisition parameters depending on the introduced fluorescent stains.
  a. Capture fluorescence on highly sensitive hybrid detectors (HyD SMD), and use a transmitted light detector for transmission images.
  b. Ensure that excitation and emission of different channels do not spectrally overlap. Bleed-through could be minimized by recording channels sequentially.
  c. For each channel, ensure sufficient signal without detector overexposure (check LUT or histogram).
  d. For the tubulin-Atto647N and DAPI stains used in this protocol, acquisition parameters were as following: tubulin-Atto647N was excited using a 638 nm diode laser and emission was captured between 642 and 782 nm; DAPI was imaged simultaneously since it does not spectrally overlap with Atto647N, excited with a 405 nm diode laser and emission was captured between 409 and 505 nm. Transmission images were also captured simultaneously. Images were acquired with 512×512 or 1024×1024 pixels, 200 Hz line scan speed and 4-line averaging.
80. Focus the sample using transmission light and the eyepiece. **NOTE:** Fluorescent stains may be visualized via the eyepiece using the fluorescence lamp, but red-emitting dyes such as Atto647N will barely be visible.
81. Select a field of view.
82. Enter live mode to view the sample on the HyD detectors and verify image acquisition parameters (step 79d). <u>Troubleshooting 7</u>.
83. Define the type of image acquisition, which is selected based on the research question.
  a. For capturing a z-stack, define the top of the cells (closest to the glass coverslip) as the starting point, and the bottom of the cells (attached to the PAAm gel) as the endpoint. Customize the step size, but keep it above 500 nm due to the axial resolution limit in confocal microscopy. We used a step size of 2 µm to visualize multiple KM12L4a cell clumps from top to bottom in 1 field of view.
  b. For more detailed visualization of protein structures, adjust the image zoom, recommended not to be lower than 4. Typically, zooms of 3 are used in our images.
  c. Z-stacking and zooming can be combined.
84. Acquire images. <u>Troubleshooting 6</u>.
85. Visualize data in LasX 3D viewer software.
  a. Open data in the software.
  b. Colors, image brightness and contrast, and gamma can be adjusted for each channel.
  c. Visualize a z-slice or zoom in 2D by selecting the desired z-plane and adding a scale bar (Fig. 7A).

**Figure 7:**
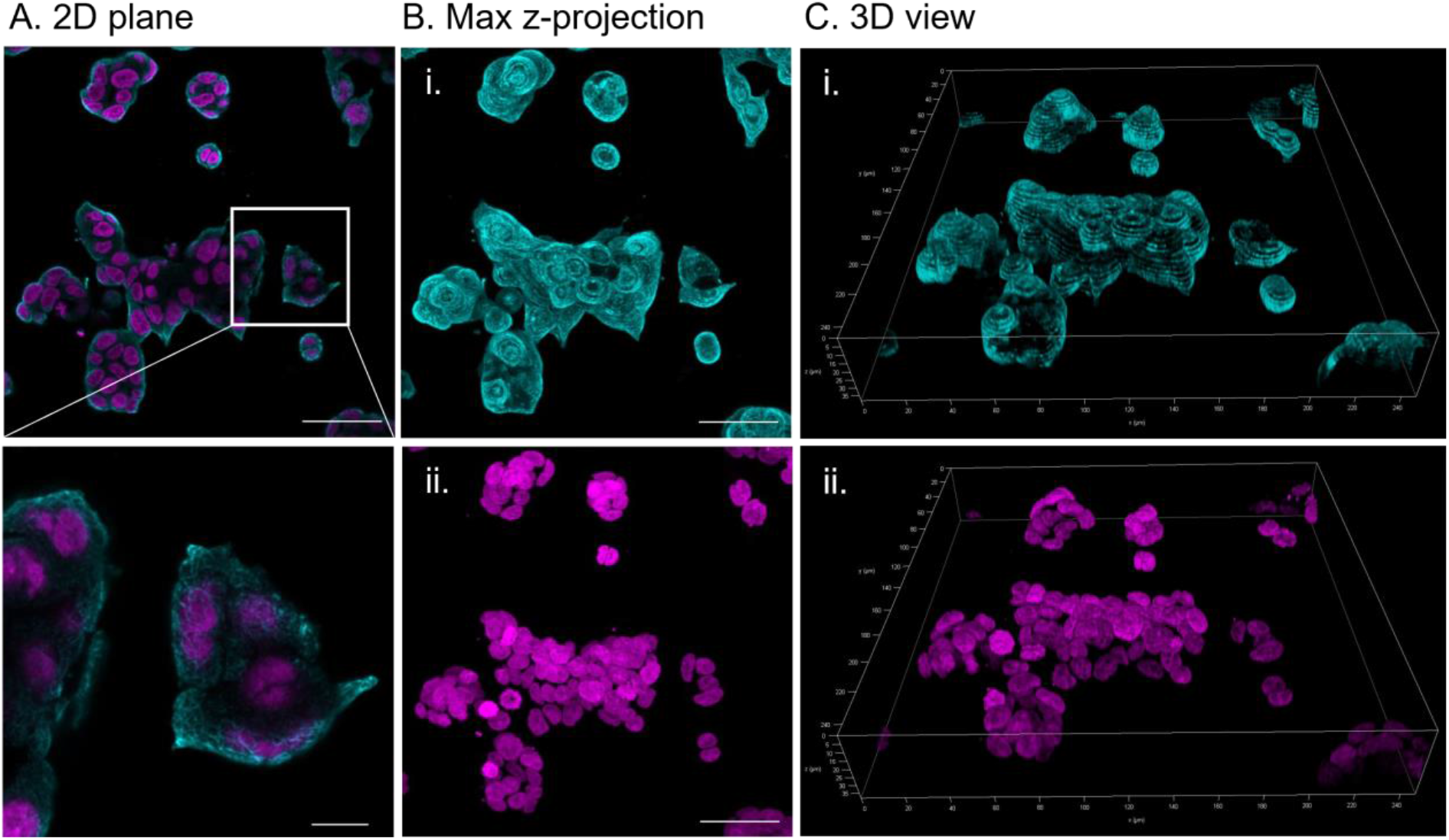
Immunolabeling and fluorescence imaging of targets on PAAm gels. **A:** One axial plane from a z-stack where both tubulin (cyan) and DAPI (magenta) are visualized. Cfr. Step 85c. Scale bar 50 µm, scale bar of zoom 10 µm. **B:** Maximum z projections of tubulin (i, cyan) and DAPI (ii, magenta). Cfr. Step 85d. Scale bars 50 µm. **C:** 3D view of tubulin (i, cyan) and DAPI (ii, magenta). Cfr. Step 85e.
  d. Alternatively, make a maximum projection of a z-stack and add a scale bar (Fig. 7B).
  e. Alternatively, visualize a z-stack in the 3D LasX viewer (Fig. 7C).
    i. Open the 3D viewer.
    ii. Set the background to black instead of grey by ticking the appropriate box.
    iii. Rotate the image by clicking and dragging the left mouse button.
    iv. Move the image by clicking and dragging the left mouse button while holding shift.
    v. Zoom by scrolling the mouse wheel.
    vi. Add a scale bar.
    vii. Add a 3D orientation inset by ticking the box “draw frame”.

## Expected outcomes

### Production of a large, unattached PAAm hydrogel

Major step 1 produces customized unattached PAAm hydrogels, swollen overnight. We typically create seven 20 mm circular gels for rheological characterization (Fig. 2D) or one 75 mm circular gel for large-batch cell culture applications (Fig. 2C), both approximately 1.2 – 1.5 mm thick. Note that gels in the pictures are shown before swelling, and that restamping is necessary after overnight swelling to obtain the final dimensions.

### Rheological characterization

Major Option A assesses the mechanical properties of PAAm hydrogels through rheology. Table 1 provides a summary of the composition and stiffness of different gels. Additional measurement details are listed in Table S1. Outcomes for the obtained key parameters, including hydrogel stiffness, are discussed below.

- Since gels are already polymerized prior to the rheology measurement and their mechanical properties should not change over short timescales, time sweep results should appear as horizontal lines (Fig. 5A). We find that the storage modulus (*G*′) is approximately 1 order of magnitude higher compared to the loss modulus (*G*′′) for the gel compositions tested here.
- When evaluating hydrogel stiffness at 1% strain, the PAAm library exhibits a broad stiffness range from 0.90 to 56.13 kPa, recapitulating both physiologically relevant conditions and diseases characterized by altered matrix stiffness.
  ∘ In agreement with previous reports, we find that increasing the concentration of either AAm monomer or MBAA crosslinker leads to an increased stiffness.^9,15^ For example, 10% AAm and 0.2% MBAA yield a stiffness of 10.07 ± 0.16 kPa (Table 1, composition 8). Higher concentrations, such as 12% AAm or 0.4% MBAA, result in increased stiffness (12.08 ± 1.95 kPa and 14.46 ± 0.06 kPa, respectively, Table 1, compositions 9 and 12). As expected, increasing both AAm and MBAA concentrations to 12% and 0.4% respectively, generates an even stiffer gel of 23.56 ± 3.58 kPa (Table 1, composition 13). However, note that extremely aberrant concentrations of either component will result in drastically altered stiffnesses, reflecting a complex relationship that has also been described in the literature.^5^ For example in our measurements, when maintaining MBAA concentration at 0.4% but increasing AAm concentrations from 10% to 12% and 18%, stiffness increases drastically from 14.46 ± 0.06 kPa to 23.56 ± 3.58 kPa and 36.70 ± 4.30 kPa respectively (Table 1, compositions 9, 13 and 14). Similarly, lowering MBAA concentrations to 0.06% significantly reduces stiffness to 0.90 ± 0.11 kPa compared to 3.73 ± 0.31 kPa for a gel with comparable AAm and 0.23% MBAA concentration (Table 1, compositions 4 and 5).
  ∘ Technical replicates obtained by stamping different areas from the same large gel demonstrate that our rheological measurements are robust and repeatable. The average error between technical replicates was 12%, while the maximum error of all compositions was found to be 18.45% (Table S1, for 4.49% AAm and 0.10% MBAA).
- Frequency sweeps are expected to appear approximately as horizontal lines, with slightly increasing storage moduli (*G*′) as the angular frequency increases (Fig. 5B). Conversely, loss moduli (*G*′′) decrease slightly with increasing angular frequency. These results are in line with previous reports.^1^
- Amplitude sweeps reveal that storage moduli (*G*′) are nearly independent of the applied oscillation strain for softer PAAm gels (here ≤ 20 kPa) (Fig. 5C). In contrast, stiffer hydrogels (here > 20 kPa) experience a decline in storage moduli when oscillation strains are increased, a trend that is more prominent for increasingly stiff gels. The effect of oscillation strain on loss moduli (*G*′′) also depends on hydrogel stiffness; softer gels exhibit less dependency on oscillation strain, while stiffer gels show an increase in loss modulus with higher oscillation strains. This behavior can be attributed to slipping of the geometry on the surface of stiffer gels when oscillation strains are high, emphasizing the importance of performing stiffness measurements at strains ≤ 1%. Note that these observations are very clear when the data is plotted in standard scale, and that these differences are less prominent when represented on the logscale.

### Large-batch cell culture & downstream applications

Major option B results in adhesion and growth of KM12L4a cells on top of functionalized PAAm hydrogels of different stiffnesses (Fig. 6). Note that KM12L4a cells are mechanosensitive, since their morphology and proliferation is influenced by the stiffness of the substrate.

Following growth, cells can be harvested for downstream applications such as Western blot, omics analysis and *in vivo* experiments (Downstream application 1). When desired, cells can be starved in medium lacking FBS, to simultaneously harvest conditioned medium (secretome) and several million cells. Note that the number of harvested cells depends on the initial seeding concentration and growth time, but possibly also on the stiffness of the gel since it affects cell proliferation.

Alternatively to cell harvesting, proteins of interest can be visualized by immunolabeling and fluorescence microscopy (Downstream application 2). Zoom and z-stack acquisitions allow detailed reconstruction of subcellular protein expression and localization in 3D (Fig. 7).

## Limitations

As mentioned earlier, large variations exist between stiffnesses reported for the same PAAm gel composition. The stiffnesses reported here are valid only when the protocol is meticulously followed, considering critical parameters like gelation temperature, the storage time of reagents or PAAm gels, and the gelation and rheology measurement temperature. To eliminate user-induced variations or when using reagents from different sources or storage conditions, we recommend independently characterizing mechanical properties of the prepared gels.

Before comparing obtained results to those reported in literature, we recommend carefully evaluating the gel polymerization conditions, as well as the measurement technique. Especially for measurement techniques at different scales, we could notice considerable differences between our own results and those reported in the literature.

Despite our best efforts to match conditions between rheology measurements and cell culture, we acknowledge that differences between the stiffness measured by rheology and the actual stiffness sensed by the cell may still arise.

- Since DPBS has a slightly higher ionic strength than DMEM, the stiffness of gels in cell culture medium will be slightly higher compared to the rheology values. Note that addition of FBS to DMEM, to create the standard cell culture medium, will not further change the stiffnesses as no salts are added.^9^
- In addition, cells will deposit their own extracellular matrix that will change not only the local stiffness but also extracellular adhesion ligands that can interact with the cellular adhesion molecules, especially when cells are cultured on a PAAm gel for longer times. However, we may argue that for the KM12L4a cell line used in this protocol, we could continue to observe changes in cell morphology for cells seeded on top of functionalized PAAm substrates of different stiffnesses during 72 h of culture.

## Troubleshooting

### Troubleshooting 1: Gel rupture immediately after polymerization

Gel ruptures or breaks into granules upon opening of the polymerization chamber or attempting to retrieve the gel from the polymerization chamber (cfr. step 10).

Potential solution: Prolonged incubation times during polymerization may cause the gel to dehydrate and make it prone to rupturing. Especially stiffer gels (> 20 kPa) may polymerize swiftly. Monitor closely the completion of the polymerization via the remaining mixture in the conical tube, add dH_2_O to the edge of the polymerization chamber as soon as polymerization is complete to avoid dehydration.

### Troubleshooting 2: Gel rupture upon handling

Especially softer gels (≤ 20 kPa) are fragile and prone to rupture (cfr. step 11 and other steps throughout the protocol when transferring gels).

Potential solutions:

- Do not attempt to scoop up the gel as a whole. Instead, gently place the scoop/spatula under the edge of the gel and transfer by sliding.
- When adding liquids with a pipet, stay away from the gel as far as possible.
- When removing liquids with a pipet, stay away from the gel as far as possible. Avoid the gel coming near the pipet tip since it may be partially sucked up. Monitor location of the gel during pipetting, as the gel might move towards the pipet tip because it is free-floating.
- When removing liquids with a pipet, remove most of the liquid with a large disposable plastic pipette (e.g. 10 mL). However, remove the final milliliters carefully and slowly using a 1 mL micropipette.

### Troubleshooting 3: Tilted gel surface

Surface of the polymerized gel appears titled instead of horizontal (cfr. step 12).

Potential solutions:

- Verify apparent tilt by placing the gel on a flat surface and ensure 100% contact between the gel mixture and the base dish, with no air bubbles in between.
- Repeat gel production and ensure polymerization is performed on a perfectly horizontal surface. Verify the Petri dish body is standing horizontally and centered in the polymerization chamber (cfr. step 9b).
- In case the problem persists, optimization of the volume for PAAm gel polymerization may be required (cfr. step 8).

### Troubleshooting 4: Inaccurate sandpaper attachment

Sandpaper attached on rheometer geometry and lower plate do not align exactly (cfr. step 19g).

Potential solution: Lower the geometry once more to check sandpaper positioning on the lower plate before pressing it down firmly (cfr. 19fvi). When attached only loosely, the sandpaper on the lower plate can still be shifted slightly.

### Troubleshooting 5: Extending the protocol to other cell types

This protocol describes steps for culturing metastatic KM12L4a colorectal cancer cells on top of collagen type I functionalized PAAm gels (Major option B). We have also used this protocol for culturing other colorectal cancer cell lines, and anticipate no issues in extending it to other adherent mammalian cell lines. If a specific cell line does not adhere to the gel, other molecules for functionalization of the gel can be used (e.g. fibrin, laminin). In addition, cell seeding concentration may need to be adjusted. Note that, while it is generally expected that cells adhere faster and spread more on stiffer substrates, mechanosensing is cell-type dependent and not all cell types seem to be mechanosensitive.^16^

### Troubleshooting 6: Immunolabeled gel floats inside imaging chamber

The immunolabeled gel moves or floats around in the 6-well glass-bottom plate (cfr. steps 75 and 84).

Potential solution: Since the gel is inverted and cells are now facing the glass-bottom surface with the gel on top, do not attempt to move or press down the gel with the round-tipped tweezers to prevent disruption of the sample. Instead, add an additional 1 or 2 mL DPBS 1x until it surrounds the gel. Then, again remove a portion to prevent sample floating.

### Troubleshooting 7: No signal in fluorescence microscopy

There is no signal visible when attempting to visualize the immunolabeled gel on the fluorescence microscope (cfr. step 82).

Potential solution:

- Verify that lasers are reaching the sample by carefully investigating the microscope from a distance after ensuring that the laser power is not set too high.
- Ensure the sample is in focus by checking transmission in the eyepieces (cfr. step 80) and in the live mode in the software.
- It is likely that DAPI signal is visible since this small molecule usually penetrates the nucleus easily, while signal from the immunolabeled target may be weak or absent.
  ∘ Verify whether the protein should be present in the cells. Use for example the online Human Protein Atlas tool (https://www.proteinatlas.org/), or previous publications.
  ∘ Verify that the primary antibody is reactive to the origin of the target (e.g. when immunolabeling a protein inside a human KM12L4a cell line, the primary antibody needs to be reactive against human). In addition, ensure that the secondary antibody is compatible with the primary antibody (e.g. for a primary antibody produced in mouse, the secondary antibody should be reactive against mouse, so for example goat-anti-mouse).
  ∘ Increase permeabilization time, or primary and secondary antibody concentrations, incubation times or temperatures. Primary antibody is frequently used at 1/200 dilutions and incubated overnight at 4 °C. Secondary antibody is frequently used at 1/1000 or 1/500 dilutions and incubated for 1-2 hours at room temperature. Do not extend secondary antibody incubations longer than 2 hours to avoid increase of non-specific binding. Use of far-red dyes is recommended since longer excitation wavelengths penetrate better through the sample.

## Resource availability

### Lead contact

Further information and requests for resources and reagents should be directed to and will be fulfilled by the lead contact, Paul Kouwer or Susana Rocha.

## Acknowledgments

The authors would like to thank colleagues from KU Leuven MIP division, especially from the group of Prof. Rocha, for their input and critical questions. We thank Fidler’s lab (MD Anderson Cancer Center) for sharing KM12 model cell lines. This work was funded by the Research Foundation - Flanders (C.C. and S.A. are both recipients of a PhD fellowship for fundamental research, FWO grant numbers 1121223N and 1S95123N). The financial support of PI20CIII/00019 and PI23CIII/00027 grants from the AES-ISCIII program cofounded by FEDER funds to R.B., and the financial support of grant PID2022-140307OB-I00 funded by MCIN/AEI/10.13039/501100011033 and by “ERDF A way of making Europe” are also gratefully acknowledged.

## Author contributions

C.C. and S.A. optimized the workflow for the synthesis of PAAm hydrogels and their functionalization. Rheological characterization was performed by C.C., with guidance from B.S. and L.G., and under supervision of P.K. A.M.-C. and R.B. assisted with optimizations for harvesting of secretome and cells. S.R. and C.C. designed the experiments and wrote the final manuscript, with graphical representations by S.A. and input from all the co-authors.

