## Supplementary material for "Synthesis and mechanical characterization of polyacrylamide (PAAm) hydrogels with different stiffnesses for large-batch cell culture applications": Table S1

Table S1: Measurement details of PAAm hydrogel library characterization. Data is organized for increasing AAm concentration, MBAA concentration is the second criterium. In datafile name, "gel" represents different technical replicates, while "test" denotes a different rheology test on the same technical replicate. Initial gap and axial force are read out when the geometry touches the PAAm gel everywhere and axial force is no longer negligible. Final gap and axial force are read out before zeroing of the axial force and starting the measurement. Storage (G') and loss (G'') moduli for time sweep at 1 % strain are given. Average and % errors on Young's moduli are calculated as described in the protocol.

| Composition (% AAm + % MBAA) | Datafile name | Initial gap (nm) | Initial axial force (N) | Final gap (nm) | Final axial force (N) | % compression | G' (Pa) | G'' (Pa) | Young's modules (E) (kPa) | Average Young's modulus (E) (kPa) | % error Young's modulus (kPa) |
|---|---|---|---|---|---|---|---|---|---|---|---|
| 4.00 + 0.17 | 20230322_4.00AAm_0.17MBAA_made 0321 18h52_gel01_test01 | 1600 | 0.02 | 1552 | 0.06 | 3.00 | 314.28 | 28.25 | 0.94 | 0.93 | 3.42 |
| | 20230322_4.00AAm_0.17MBAA_made 0321 18h52_gel02_test01 | 1680 | 0.02 | 1630 | 0.04 | 2.98 | 319.19 | 27.08 | 0.96 | | |
| | 20230322_4.00AAm_0.17MBAA_made 0321 18h52_gel03_test01 | 1400 | 0.01 | 1340 | 0.05 | 4.29 | 298.32 | 22.43 | 0.89 | | |
| 4.49 + 0.10 | 20230322_4.94AAm_0.10MBAA_made 0321 19h10_gel01_test01 | 1550 | 0.02 | 1503 | 0.07 | 3.03 | 339.32 | 25.79 | 1.02 | 1.23 | 18.45 |
| | 20230322_4.94AAm_0.10MBAA_made 0321 19h10_gel01_test02 | 1470 | 0.02 | 1400 | 0.17 | 4.76 | 412.89 | 29.99 | 1.24 | | |
| | 20230322_4.94AAm_0.10MBAA_made 0321 19h10_gel02_test01 | 1650 | 0.01 | 1590 | 0.08 | 3.64 | 484.48 | 36.60 | 1.45 | | |
| | 20230322_4.94AAm_0.10MBAA_made 0321 19h10_gel03_test01 | 1650 | 0.02 | 1600 | 0.06 | 3.03 | 290.77 | 25.54 | 0.87 | | |
| | 20230322_4.94AAm_0.10MBAA_made 0321 19h10_gel03_test02 | ND | ND | 1400 | 0.30 | ND | 519.52 | 41.73 | 1.56 | | |
| 5.00 + 0.15 | 20230322_5.00AAm_0.15MBAA_made 0321 19h13_gel01_test01 | 1400 | 0.03 | 1358 | 0.11 | 3.00 | 544.62 | 45.48 | 1.63 | 2.19 | 15.71 |
| | 20230322_5.00AAm_0.15MBAA_made 0321 19h13_gel01_test02 | 1300 | 0.25 | 1250 | 0.37 | 3.85 | 839.58 | 60.42 | 2.52 | | |
| | 20230322_5.00AAm_0.15MBAA_made 0321 19h13_gel02_test01 | 1420 | 0.04 | 1377 | 0.18 | 3.03 | 686.05 | 56.16 | 2.06 | | |

| | | | | | | | | | | | |
|---|---|---|---|---|---|---|---|---|---|---|---|
| | 20230322_5.00AAm_0.15MBAA_made 0321 19h13_gel02_test02 | ND | ND | 1270 | 0.45 | ND | 900.67 | 67.29 | 2.70 | | |
| | 20230322_5.00AAm_0.15MBAA_made 0321 19h13_gel03_test01 | 1580 | 0.07 | 1532 | 0.22 | 3.04 | 678.62 | 83.30 | 2.04 | | |
| 5.50 + 0.23 | 20230322_5.50AAm_0.23MBAA_made 0321 18h46_gel01_test01 | 1500 | 0.02 | 1450 | 0.11 | 3.33 | 1163.78 | 79.43 | 3.49 | 3.73 | 8.39 |
| | 20230322_5.50AAm_0.23MBAA_made 0321 18h46_gel01_test02 | 1460 | 0.03 | 1416 | 0.20 | 3.01 | 1388.50 | 89.48 | 4.17 | | |
| | 20230322_5.50AAm_0.23MBAA_made 0321 18h46_gel02_test01 | 1450 | 0.04 | 1406 | 0.16 | 3.03 | 1284.67 | 84.02 | 3.85 | | |
| | 20230322_5.50AAm_0.23MBAA_made 0321 18h46_gel03_test02 | 1380 | 0.04 | 1338 | 0.13 | 3.04 | 1135.82 | 91.93 | 3.41 | | |
| 6.00 + 0.06 | 20230321_6.00AAm_0.06MBAA_made0320 12h56_gel01_test02 | 1800 | 0.03 | 1750 | 0.06 | 2.78 | 269.95 | 13.37 | 0.81 | 0.90 | 11.65 |
| | 20230321_6.00AAm_0.06MBAA_made0320 12h56_gel02_test01 | 1850 | 0.04 | 1800 | 0.07 | 2.70 | 286.13 | 20.55 | 0.86 | | |
| | 20230321_6.00AAm_0.06MBAA_made0320 12h56_gel03_test01 | 1900 | 0.02 | 1830 | 0.07 | 3.68 | 346.58 | 17.85 | 1.04 | | |
| 7.50 + 0.32 | 20230322_7.50AAm_0.32MBAA_made 0321 18h17_gel01_test01 | 1550 | 0.04 | 1500 | 0.18 | 3.23 | 2378.91 | 170.53 | 7.14 | 7.45 | 3.75 |
| | 20230322_7.50AAm_0.32MBAA_made 0321 18h17_gel02_test01 | ND | ND | 1500 | 0.20 | ND | 2502.88 | 173.84 | 7.51 | | |
| | 20230322_7.50AAm_0.32MBAA_made 0321 18h17_gel03_test01 | 1580 | 0.15 | 1530 | 0.23 | 3.16 | 2568.65 | 182.24 | 7.71 | | |
| 9.00 + 0.14 | 20230322_9.00AAm_0.14MBAA_made 0321 18h13_gel01_test01 | 1900 | 0.04 | 1843 | 0.18 | 3.00 | 1758.46 | 101.98 | 5.28 | 5.91 | 9.05 |
| | 20230322_9.00AAm_0.14MBAA_made 0321 18h13_gel02_test01 | 1750 | 0.06 | 1680 | 0.20 | 4.00 | 1853.42 | 101.73 | 5.56 | | |
| | 20230322_9.00AAm_0.14MBAA_made 0321 18h13_gel02_test02 | 1700 | 0.08 | 1630 | 0.18 | 4.12 | 2074.21 | 107.93 | 6.22 | | |
| | 20230322_9.00AAm_0.14MBAA_made 0321 18h13_gel03_test01 | ND | ND | 1770 | 0.18 | ND | 2190.87 | 118.49 | 6.57 | | |
| 10.00 + 0.20 | 20230323_10.00AAm_0.20MBAA_made 0322 18h57_gel01_test01 | 1750 | 0.02 | 1680 | 0.32 | 4.00 | 3752.15 | 303.48 | 11.26 | 10.07 | 15.97 |
| | 20230323_10.00AAm_0.20MBAA_made 0322 18h57_gel02_test01 | 1600 | 0.02 | 1520 | 0.25 | 5.00 | 2667.06 | 208.71 | 8.00 | | |
| | 20230323_10.00AAm_0.20MBAA_made 0322 18h57_gel03_test01 | 1800 | 0.04 | 1720 | 0.34 | 4.44 | 3650.55 | 283.82 | 10.95 | | |
| 10.00 + 0.40 | 20230323_10.00AAm_0.40MBAA_made 0322 18h53_gel01_test01 | 1620 | 0.05 | 1570 | 0.47 | 3.09 | 5178.96 | 489.43 | 15.54 | 14.46 | 6.13 |
| | 20230323_10.00AAm_0.40MBAA_made 0322 18h53_gel02_test01 | ND | ND | 1400 | 0.40 | ND | 4572.01 | 415.08 | 13.72 | | |
| | 20230323_10.00AAm_0.40MBAA_made 0322 18h53_gel03_test01 | ND | ND | 1400 | 0.40 | ND | 4712.70 | 443.54 | 14.14 | | |
| 10.00 + 0.43 | 20230322_10.00AAm_0.43MBAA_made 0321 17h43_gel01_test01 | 1370 | 0.27 | 1320 | 0.20 | 3.65 | 5954.05 | 461.18 | 17.86 | 16.90 | 13.64 |

| | | | | | | | | | | | |
|---|---|---|---|---|---|---|---|---|---|---|---|
| | 20230322_10.00AAm_0.43 MBAA_made 0321 17h43_gel02_test01 | 1550 | 0.15 | 1500 | 0.38 | 3.23 | 5792.89 | 485.04 | 17.38 | | |
| | 20230322_10.00AAm_0.43 MBAA_made 0321 17h43_gel03_test01 | 1500 | 0.02 | 1455 | 0.13 | 3.00 | 4041.15 | 364.95 | 12.12 | | |
| | 20230322_10.00AAm_0.43 MBAA_made 0321 17h43_gel03_test02 | ND | ND | 1400 | 0.38 | ND | 5570.60 | 444.93 | 16.71 | | |
| | 20230322_10.00AAm_0.43 MBAA_made 0321 17h43_gel04_test01 | 1650 | 0.04 | 1600 | 0.30 | 3.03 | 5189.28 | 487.60 | 15.57 | | |
| | 20230323_10.00AAm_0.43 MBAA_made 0322 18h01_gel01_test01 | ND | ND | 1450 | 0.33 | ND | 7483.08 | 743.03 | 22.45 | | |
| | 20230323_10.00AAm_0.43 MBAA_made 0322 18h01_gel02_test01 | ND | ND | 1500 | 0.26 | ND | 4817.83 | 455.30 | 14.45 | | |
| | 20230323_10.00AAm_0.43 MBAA_made 0322 18h01_gel03_test01 | ND | ND | 1500 | 0.26 | ND | 6221.29 | 664.83 | 18.66 | | |
| 10.50 + 0.45 | 20230321_10.50AAm_0.45 MBAA_made0320 12h10_gel02_test01 | 1900 | 0.15 | 1850 | 0.60 | 2.63 | 8890.35 | 880.13 | 26.67 | 20.95 | 14.04 |
| | 20230321_10.50AAm_0.45 MBAA_made0320 12h10_gel03_test01 | 1700 | 0.20 | 1650 | 0.55 | 2.94 | 5813.23 | 498.14 | 17.44 | | |
| | 20230321_10.50AAm_0.45 MBAA_made0320 12h10_gel03_test02 | 1680 | 0.19 | 1600 | 1.00 | 4.76 | 7485.46 | 606.91 | 22.46 | | |
| | 20230323_10.50AAm_0.45 MBAA_made 0322 18h03_gel02_test01 | 1600 | 0.06 | 1520 | 0.49 | 5.00 | 6040.03 | 518.23 | 18.12 | | |
| | 20230323_10.50AAm_0.45 MBAA_made 0322 18h03_gel03_test01 | 1550 | 0.03 | 1460 | 0.41 | 5.81 | 6688.24 | 631.08 | 20.06 | | |
| 12.00 + 0.20 | 20230322_12.00AAm_0.20 MBAA_made 0321 18h38_gel01_test01 | 1980 | 0.03 | 1920 | 0.27 | 3.03 | 4578.30 | 305.09 | 13.73 | 12.08 | 16.14 |
| | 20230322_12.00AAm_0.20 MBAA_made 0321 18h38_gel02_test01 | 1700 | 0.07 | 1650 | 0.23 | 2.94 | 4153.10 | 258.00 | 12.46 | | |
| | 20230322_12.00AAm_0.20 MBAA_made 0321 18h38_gel03_test01 | 1800 | 0.02 | 1746 | 0.20 | 3.00 | 4483.87 | 306.23 | 13.45 | | |
| | 20230323_12.00AAm_0.20 MBAA_made 0322 18h07_gel01_test01 | 1820 | 0.03 | 1730 | 0.27 | 4.95 | 3501.10 | 222.09 | 10.50 | | |
| | 20230323_12.00AAm_0.20 MBAA_made 0322 18h07_gel02_test02 | 1880 | 0.08 | 1820 | 0.35 | 3.19 | 6351.21 | 470.24 | 19.05 | | |
| | 20230323_12.00AAm_0.20 MBAA_made 0322 18h07_gel03_test01 | 1700 | 0.06 | 1650 | 0.25 | 2.94 | 3419.05 | 263.94 | 10.26 | | |
| | 20230323_12.00AAm_0.20 MBAA_made 0322 18h07_gel04_test01 | 1600 | 0.06 | 1550 | 0.25 | 3.13 | 4470.82 | 279.44 | 13.41 | | |
| | 20230323_12.00AAm_0.20 MBAA_made 0322 18h57_gel01_test01 | 1620 | 0.04 | 1550 | 0.24 | 4.32 | 2885.30 | 214.76 | 8.66 | | |
| | 20230323_12.00AAm_0.20 MBAA_made 0322 18h57_gel02_test01 | 1650 | 0.03 | 1570 | 0.17 | 4.85 | 3033.11 | 221.21 | 9.10 | | |
| | 20230323_12.00AAm_0.20 MBAA_made 0322 18h57_gel03_test01 | 1930 | 0.07 | 1850 | 0.26 | 4.15 | 3402.23 | 253.77 | 10.21 | | |
| 12.00 + 0.40 | 20230321_12.00AAm_0.40 MBAA_made0320 10h40_gel02_test02 | 2000 | 0.02 | 1935 | 0.67 | 3.25 | 11059.18 | 1017.51 | 33.18 | 23.56 | 15.19 |

| | | | | | | | | | | |
|---|---|---|---|---|---|---|---|---|---|---|
| | 20230321_12.00AAm_0.40 MBAA_made0320 10h40_gel03_test01 | 1650 | 0.37 | 1600 | 0.66 | 3.03 | 7414.93 | 551.53 | 22.24 | | |
| | 20230321_12.00AAm_0.40 MBAA_made0320 10h40_gel04_test02 | 1550 | 0.03 | 1500 | 0.53 | 3.23 | 6905.83 | 471.19 | 20.72 | | |
| | 20230323_12.00AAm_0.40 MBAA_made 0322 18h09_gel01_test01 | 1750 | 0.02 | 1650 | 0.46 | 5.71 | 6968.78 | 587.46 | 20.91 | | |
| | 20230323_12.00AAm_0.40 MBAA_made 0322 18h09_gel02_test02 | 1890 | 0.07 | 1820 | 0.35 | 3.70 | 10328.42 | 1083.21 | 30.99 | | |
| | 20230323_12.00AAm_0.40 MBAA_made 0322 18h09_gel03_test01 | 1600 | 0.06 | 1550 | 0.25 | 3.13 | 6418.40 | 567.30 | 19.26 | | |
| | 20230323_12.00AAm_0.40 MBAA_made 0322 18h55_gel01_test01 | 1580 | 0.08 | 1530 | 0.50 | 3.16 | 6535.16 | 582.78 | 19.61 | | |
| | 20230323_12.00AAm_0.40 MBAA_made 0322 18h55_gel02_test01 | 1730 | 0.14 | 1670 | 0.50 | 3.47 | 7915.32 | 721.44 | 23.75 | | |
| | 20230323_12.00AAm_0.40 MBAA_made 0322 18h55_gel03_test01 | 1740 | 0.20 | 1680 | 0.35 | 3.45 | 7138.08 | 691.59 | 21.41 | | |
| 18.00 + 0.40 | 20230321_18.00AAm_0.40 MBAA_made0320 12h23_gel01_test01 | 1880 | 0.02 | 1825 | 0.50 | 2.93 | 13810.28 | 1395.90 | 41.43 | 36.70 | 11.72 |
| | 20230321_18.00AAm_0.40 MBAA_made0320 12h23_gel02_test01 | 1680 | 0.07 | 1625 | 0.50 | 3.27 | 10043.60 | 720.89 | 30.13 | | |
| | 20230321_18.00AAm_0.40 MBAA_made0320 12h23_gel03_test01 | 1680 | 0.05 | 1610 | 0.50 | 4.17 | 11133.82 | 961.38 | 33.40 | | |
| | 20230322_18.00AAm_0.40 MBAA_made 0321 18h57_gel01_test01 | 1750 | 0.04 | 1680 | 0.56 | 4.00 | 11256.38 | 714.43 | 33.77 | | |
| | 20230322_18.00AAm_0.40 MBAA_made 0321 18h57_gel02_test01 | 1950 | 0.03 | 1890 | 0.50 | 3.08 | 13407.36 | 792.20 | 40.22 | | |
| | 20230322_18.00AAm_0.40 MBAA_made 0321 18h57_gel03_test01 | 2200 | 0.03 | 2135 | 0.50 | 2.95 | 13739.74 | 1070.67 | 41.22 | | |
| 23.00 + 0.45 | 20230321_23.00AAm_0.45 MBAA_made0320 12h31_gel01_test01 | 1750 | 0.06 | 1690 | 0.50 | 3.43 | 13641.76 | 1162.21 | 40.93 | 47.25 | 11.66 |
| | 20230321_23.00AAm_0.45 MBAA_made0320 12h31_gel02_test01 | 1750 | 0.01 | 1680 | 0.50 | 4.00 | 15824.20 | 1293.35 | 47.47 | | |
| | 20230321_23.00AAm_0.45 MBAA_made0320 12h31_gel03_test01 | 1750 | 0.01 | 1680 | 0.50 | 4.00 | 17786.70 | 1619.59 | 53.36 | | |
| 28.90 + 0.50 | 20230321_28.90AAm_0.50 MBAA_made0320 12h37_gel01_test01 | 1750 | 0.02 | 1680 | 0.50 | 4.00 | 19018.54 | 1423.834 | 57.06 | 56.13 | 12.14 |
| | 20230321_28.90AAm_0.50 MBAA_made0320 12h37_gel02_test01 | ND | ND | 1925 | 0.56 | ND | 21106.44 | 1534.902 | 63.32 | | |
| | 20230321_28.90AAm_0.50 MBAA_made0320 12h37_gel03_test01 | 1730 | 0.02 | 1665 | 0.44 | 3.76 | 16008.475 | 1139.1675 | 48.03 | | |
